# A genome-wide almanac of co-essential modules assigns function to uncharacterized genes

**DOI:** 10.1101/827071

**Authors:** Michael Wainberg, Roarke A. Kamber, Akshay Balsubramani, Robin M. Meyers, Nasa Sinnott-Armstrong, Daniel Hornburg, Lihua Jiang, Joanne Chan, Ruiqi Jian, Mingxin Gu, Anna Shcherbina, Michael M. Dubreuil, Kaitlyn Spees, Michael P. Snyder, Anshul Kundaje, Michael C. Bassik

**Author notes:** These authors contributed equally (Michael Wainberg, Roarke A. Kamber, Akshay Balsubramani).

## Abstract

A central remaining question in the post-genomic era is how genes interact to form biological pathways. Measurements of gene dependency across hundreds of cell lines have been used to cluster genes into ‘co-essential’ pathways, but this approach has been limited by ubiquitous false positives. Here, we develop a statistical method that enables robust identification of gene co-essentiality and yields a genome-wide set of functional modules. This almanac recapitulates diverse pathways and protein complexes and predicts the functions of 102 uncharacterized genes. Validating top predictions, we show that *TMEM189* encodes plasmanylethanolamine desaturase, the long-sought key enzyme for plasmalogen synthesis. We also show that *C15orf57* binds the AP2 complex, localizes to clathrin-coated pits, and enables efficient transferrin uptake. Finally, we provide an interactive web tool for the community to explore the results (coessentiality.net). Our results establish co-essentiality profiling as a powerful resource for biological pathway identification and discovery of novel gene functions.

## INTRODUCTION

A fundamental and still largely unresolved question in biology is how finite numbers of genes generate the vast phenotypic complexity of cells and organisms (Barabási and Oltvai, 2004; Chuang et al., 2010). With the understanding that modules of interacting genes represent a key layer of biological organization, the complete identification of such functional modules and their constituent genes has emerged as a central goal of systems biology (Costanzo et al., 2016; Hartwell et al., 1999; Horlbeck et al., 2018; Stuart et al., 2003). However, efforts to map genetic interactions and biological modules at genome scale have been hindered by the enormous number of possible gene-gene interactions: assaying all pairs of genetic interactions among the approximately 20,000 human genes (Harrow et al., 2012) would require 200 million distinct readouts. Furthermore, despite substantial progress in elucidating the functions of individual genes in recent decades through both targeted studies and unbiased approaches (Alonso and Ecker, 2006; Carpenter and Sabatini, 2004; Mohr et al., 2014; Shalem et al., 2015), hundreds of human genes remain functionally uncharacterized.

Pioneering work in yeast measured pairwise genetic interactions in high throughput by quantifying the fitness of double knockout strains (Tong, 2004; Tong et al., 2001); more recently, this work has been extended into a genome-wide map of yeast genetic interactions and modules (Costanzo et al., 2010, 2016). In human cells, which unlike yeast cannot be crossed to generate double-knockout mutants, a key advance towards genetic interaction mapping has been the development of genome-scale CRISPR/Cas9 and RNAi screens (Mohr et al., 2014; Shalem et al., 2015) which have been repurposed to perform pairs of perturbations (Bassik et al., 2013; Boettcher et al., 2018; Du et al., 2017; Han et al., 2017; Horlbeck et al., 2018; Rosenbluh et al., 2016; Shen et al., 2017). Yet despite considerable successes, double-perturbation genetic interaction mapping is inherently limited by the combinatorial explosion of gene pairs, with the largest human genetic interaction map to date (Horlbeck et al., 2018) having only assayed 222,784 gene pairs, or ∼0.1% of all possible genetic interactions, thus far precluding the generation of a genome-wide map of functional modules in human cells.

A complementary approach that circumvents this limitation is to measure the fitness of single-gene perturbations across multiple conditions, and map putative functional interactions by correlating the resulting phenotypic profiles (Figure S1A), referred to as co-essentiality mapping. Both co-essentiality mapping and genetic interaction mapping measure gene essentiality across many different genetic backgrounds, but whereas the background for genetic interaction mapping is the knockout of a single partner gene, for co-essentiality mapping it is the mutational and transcriptional profile of a cell line. Co-essentiality mapping across diverse cancer cell lines has recently been used to group genes into pathways and in some cases has identified novel gene functions (Boyle et al., 2018; Kim et al., 2019; McDonald et al., 2017; Pan et al., 2018; Rauscher et al., 2018; Wang et al., 2017).

Co-essentiality mapping, however, has its own fundamental limitation: unlike double-perturbation mapping, where each pair of gene knockouts is independent, measurements in two different cell lines may be strongly related, for instance because some pairs of cell lines are derived from the same tissue or lineage. Existing approaches fail to account for violations of independence, leading to inflated *p*-values, incorrect determinations of statistical significance, and an inability to identify gene co-essentiality relationships in a robust, systematic manner (Figure S1B). In this study, we address this critical limitation of co-essentiality mapping with a novel statistical method that explicitly accounts for the non-independence of cell lines. We apply the method to a dataset of genome-wide CRISPR screens in 485 diverse cancer cell lines (Tsherniak et al., 2017) and find significantly improved enrichment for known pathway interactions and protein complexes.

We find that these analytical advances greatly improve our ability to detect *bona fide* functional modules. We generate a genome-wide almanac of co-essential modules, which both recapitulate diverse known pathways and protein complexes and nominate putative functions for 102 poorly characterized genes. We experimentally validate two such genes: we identify *TMEM189* as the gene encoding the plasmanylethanolamine desaturase (PEDS) orphan enzyme required for synthesis of plasmalogen lipids, one of the most abundant lipid classes in the human body; and we discover a role for *C15orf57* in regulating clathrin-mediated endocytosis. Finally, to accelerate further biological discovery using this resource, we present an interactive web tool that enables visualization and analysis of co-essential gene pairs and modules.

## RESULTS

### A genome-wide map of co-essential interactions

To map co-essential interactions across compendia of genome-wide screens while accounting for non-independence of cell lines, we devised a novel approach based on generalized least squares (GLS), a classic statistical technique (Aitkin, 1935) (see Methods). We applied the approach (Figure 1A) to a dataset of CRISPR screens in 485 cell lines from the Achilles project (Tsherniak et al., 2017), with gene-level essentiality scores corrected for copy number and guide efficacy using the CERES algorithm (Meyers et al., 2017). We noted the remarkably effective statistical calibration of the method. Since the percentage of gene pairs expected to have detectable functional interactions is much less than 50% (Horlbeck et al., 2018), the median *p*-value across gene pairs ought to be very close to 0.5 for a well-calibrated method. Indeed, we found that the median GLS *p*-value was 0.48, indicating near-perfect calibration, while the median Pearson correlation *p*-value on the same dataset was 0.21, indicating substantial inflation and false-positive co-essential gene pairs (Figure 1B). We provide each gene’s significant co-essential interactors at a false discovery rate of 10% (**Table S1**).

**Figure 1:**
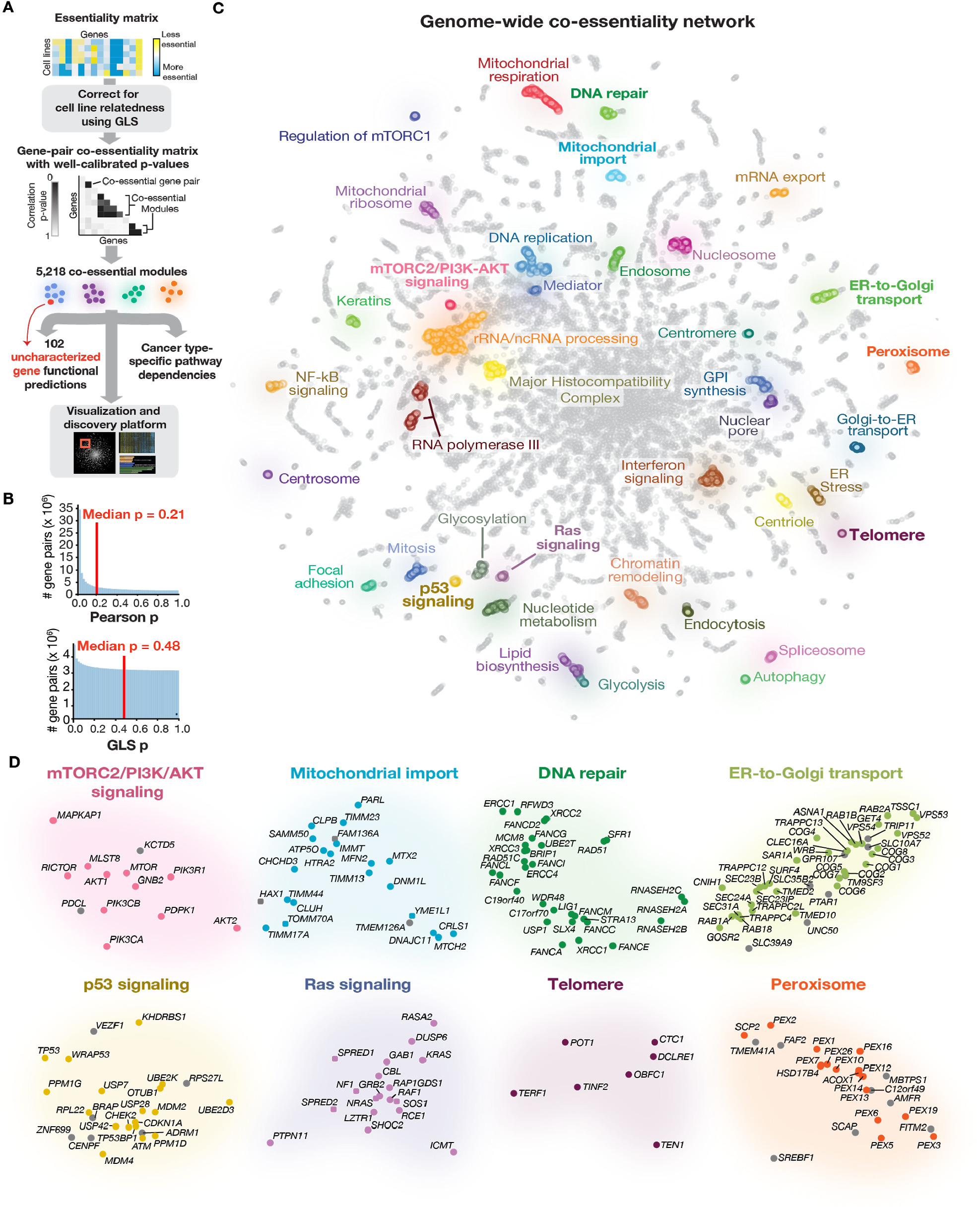
Construction of a genome-wide co-essentiality network. **(A)** Overview of our approach. **(B)** Histograms of GLS and Pearson correlations across all pairs of genes. **(C)** Global structure of the co-essentiality network, with manually annotated ‘neighborhoods’ highly enriched for particular pathways and complexes. Bolded neighborhood labels are highlighted in (D). **(D)** Selected neighborhoods, with manually-defined known pathway members indicated in color and other genes in gray. See also Figure S1.

Even while correcting for *p*-value inflation, GLS still has substantial power to detect co-essential interactions. Around 80% of genes have at least one co-essential partner at 10% FDR (Figure S2), and 40% of genes have at least ten partners: in all, we detect 93,575 significant co-essential gene pairs. 99.4% of all partners are positively correlated, with the remaining 0.6% negatively correlated. We noted that in many cases, negative correlations occur when one gene negatively regulates the other: for instance, *TP53* is negatively correlated with *MDM2* (*p* = 1 × 10^-12^), which ubiquitinates p53 to mark it for degradation (Moll and Petrenko, 2003); *HER2* is negatively correlated with *PHLDA2* (*p* = 5 × 10^-6^), which was recently shown to inhibit *HER2* signaling (Wang et al., 2018); and *MAPK1* is negatively correlated with *DUSP6* (*p* = 2 × 10^-6^), a phosphatase that inactivates several MAP kinases including MAPK1 (Furukawa et al., 2008). A second class of negative correlation arises from genes with similar functions that are active in mutually exclusive cell types, such as *MYC* and *MYCN* (*p* = 3 × 10^-11^) (Rickman et al., 2018).

**Figure 2:**
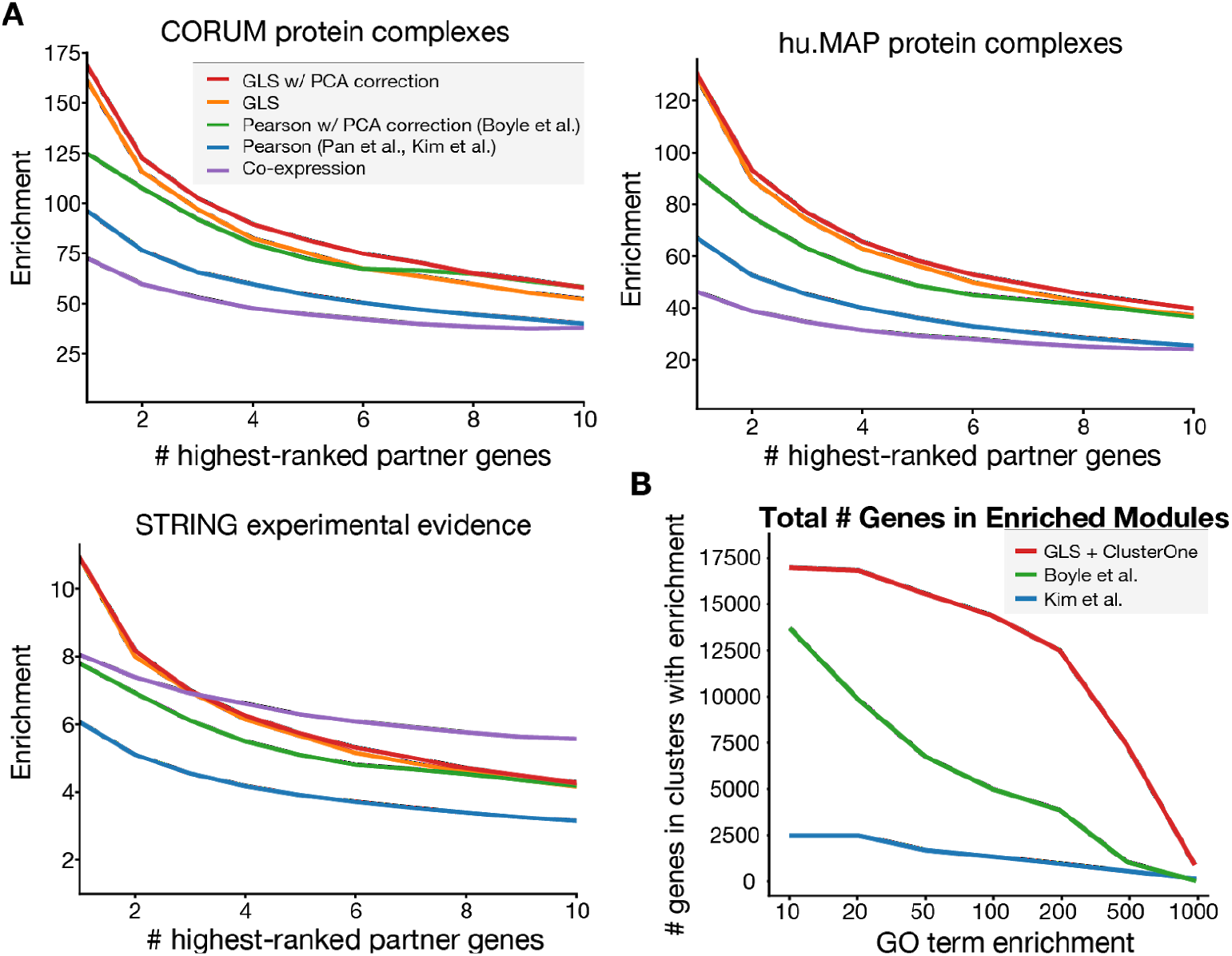
GLS improves recall of known functional interactions in co-essential gene pairs and modules. **(A)** Enrichment of interactions from GLS- and Pearson-based co-essentiality (with/without PCA-based bias correction) using the DepMap dataset, as well as co-expression using the COXPRESdb dataset, in CORUM, hu.MAP and STRING, considering the top 1-10 partners per gene. **(B)** Number of genes in clusters/modules at least N-fold enriched for some GO term, excluding the gene itself from the enrichment calculation, for various N from 10 to 1000. See also Figures S2, S3, S6 and S7

Crucially, even though more essential genes tend to have more partners, 70% of the 10% least essential genes have at least one partner at 10% FDR, and nearly half of these least essential genes have at least one partner at 1% FDR (Figure S2). This suggests that, rather than being limited to detecting interactions among only strongly essential genes, the focus of previous co-essentiality mapping efforts (Kim et al., 2019), co-essentiality is a genome-wide tool for pathway mapping.

We developed a method to visualize genes in a genome-wide interaction map based on their co-essentiality profiles by placing more strongly co-essential gene pairs closer together, inspired by similar visualizations based on yeast genetic interaction maps (Costanzo et al., 2010, 2016). We found that naive application of dimensionality reduction techniques such as Principal Component Analysis (PCA) and Uniform Manifold Approximation and Projection (UMAP) (McInnes et al., 2018) failed to effectively expose functional relationships between genes due to difficulty modeling the multi-scale nature of the co-essentiality network; previous attempts at visualizing co-essentiality networks (e.g. McDonald et al., 2017) also suffer from a similar lack of discernible structure. Instead, we first applied diffusion maps (Coifman and Lafon, 2006), a technique from spectral graph theory, to separate coarse- and fine-scale components before applying UMAP (see Methods). To further improve the layout, we incorporated module membership (defined below) into the diffusion map in addition to pairwise co-essentiality. To showcase the power of this approach, we manually annotated 39 ‘neighborhoods’ within the interaction map highly enriched for a particular pathway or complex (Figure 1C, D); collectively, these pathways and complexes encompass many of the major aspects of cell biology.

### Co-essentiality complements co-expression in mapping biological pathways

We next investigated whether the improved calibration of GLS translated into improved prioritization of co-functional gene pairs. To do this, we used an established benchmarking strategy (Pan et al., 2018) to measure how accurately GLS could recall the top 1 to 10 interaction partners of each gene when compared with Pearson correlation. The performance was measured using three databases of interactors previously benchmarked in Pan et al.: CORUM, a manually curated protein complex database (Ruepp et al., 2008); hu.MAP, a database of protein-protein interactions detected by mass spectrometry experiments (Drew et al., 2017); and STRING, a database of co-functional interactions integrating multiple sources of direct and indirect evidence (Szklarczyk et al., 2017). We found that GLS consistently prioritized genes more effectively than several other methods, including Pearson correlation bias-corrected with PCA using olfactory receptor genes as a gold-standard negative set (Boyle et al., 2018), across all three databases and across a wide variety of rank thresholds (Figure 2A). For instance, the top-ranked partners for each gene are approximately 160-fold enriched for CORUM interactions for GLS compared to 120-fold for bias-corrected Pearson correlation; for hu.MAP, 130-fold versus 90-fold enriched; and for STRING, 7.5-fold versus 5.5-fold enriched. Remarkably, failing to perform PCA-based bias correction significantly degrades the performance of Pearson correlation but not GLS, suggesting that GLS is able to automatically perform bias correction without requiring a putatively non-essential gene set like olfactory receptors.

We also compared co-essentiality to co-expression, a complementary approach to assessing co-functionality, using the COXPRESdb database (Okamura et al., 2015). We observed that co-essentiality substantially outperformed co-expression in recall of protein complexes and physical interactions recorded in the CORUM and hu.MAP databases, but performance was more equivocal for STRING (Figure 2A), with co-essentiality outperforming co-expression only for top-ranked partner genes. Of note, STRING integrates seven sources of evidence (experimental evidence, other pathway/complex databases, co-expression, literature text-mining, genomic co-localization across species, co-occurrence across species, and existence of a gene-gene fusion in any species); to reduce the potential for circularity, we restricted to gene pairs supported by experimental evidence. Collectively, these results suggest that co-essentiality and co-expression may have complementary roles in biological pathway mapping, with co-essentiality being better-suited for detecting protein complexes and direct physical interactions and co-expression being better-suited for detecting more indirect functional relationships such as regulatory relationships.

Co-essentiality was particularly effective in detecting interactions for a number of key cancer drivers. For example, 8 of *TP53*’s 10 significant co-essential partners are known interactors (*USP28*, *CDKN1A*, *TP53BP1*, *MDM2*, *CHEK2*, *ATM*, *PPM1D*, *UBE2K*) compared with only 3 of the top 10 co-expressed partners in COXPRESdb (Table S2). For *KRAS*, 3 of 5 significant co-essential partners are known interactors compared to none of the top 5 co-expressed partners; and for *BRCA1*, 3 of 6 co-essential partners are known interactors compared to 1 of 6 for co-expression.

### Co-essential modules recapitulate known pathways and nominate novel members

To group genes into modules based on their co-essentiality profiles from GLS, we used ClusterONE (Nepusz et al., 2012), a commonly-used algorithm originally developed for the *de novo* discovery of protein complexes from protein-protein interaction data (see Methods). Crucially, the modules generated by ClusterONE are allowed to be overlapping, enabling pleiotropic genes to be constituents of multiple modules. One major parameter that affects the quality of ClusterONE module detection is the module density *d*, which determines (on a 0-to-1 scale) how strong the internal connections within a cluster must be relative to the connections on the edge of the cluster between members and non-members. It has previously been observed that inferring networks at multiple scales helps provide the most complete picture of biological systems (Dutkowski et al., 2013; Kramer et al., 2014); accordingly, we found that lower values of *d* (e.g. *d* = 0.2) led to larger modules and better performance on STRING, while larger values (e.g. d = 0.9) led to smaller modules and better performance at recapitulating complexes and physical interactions from CORUM and hu.MAP, with intermediate values (e.g. *d* = 0.5) striking a balance between the two (Figure S3). Because modules generated with different values of *d* capture different types of biological pathways, we generated a combined list of modules using *d* = 0.2 (N = 168), *d* = 0.5 (N = 1892) and *d* = 0.9 (N = 3169) (**Table S3**).

The 5,218 co-essential modules in this almanac, containing between 4 and 741 genes, correspond to a wide range of biological pathways (**Table S3**). To estimate the fraction of the genome our modules assign a putative function, we counted the number of genes included in a module that is highly (at least 100-fold) enriched for some GO term. By this metric, our set of co-essential modules assign putative functions to 14,383 genes, a much larger fraction of the genome compared to previous approaches used to cluster genes based on co-essentiality profiles (Figure 2B).

Among the 1,311 modules with greater than 100-fold enrichments are modules highly enriched for genes involved in growth regulation (Figures 3A, B), autophagy (Figure 3C), cell-cell signaling (Figure 3D), the DNA damage response (Figure 3E), innate immunity (Figure 3F), glycolysis (Figure 3G), transcriptional regulation (Figures 3H, I), the cell cycle (Figure 3J), and mitochondrial respiration (Figure 3K), among many others (**Table S3**).

**Figure 3:**
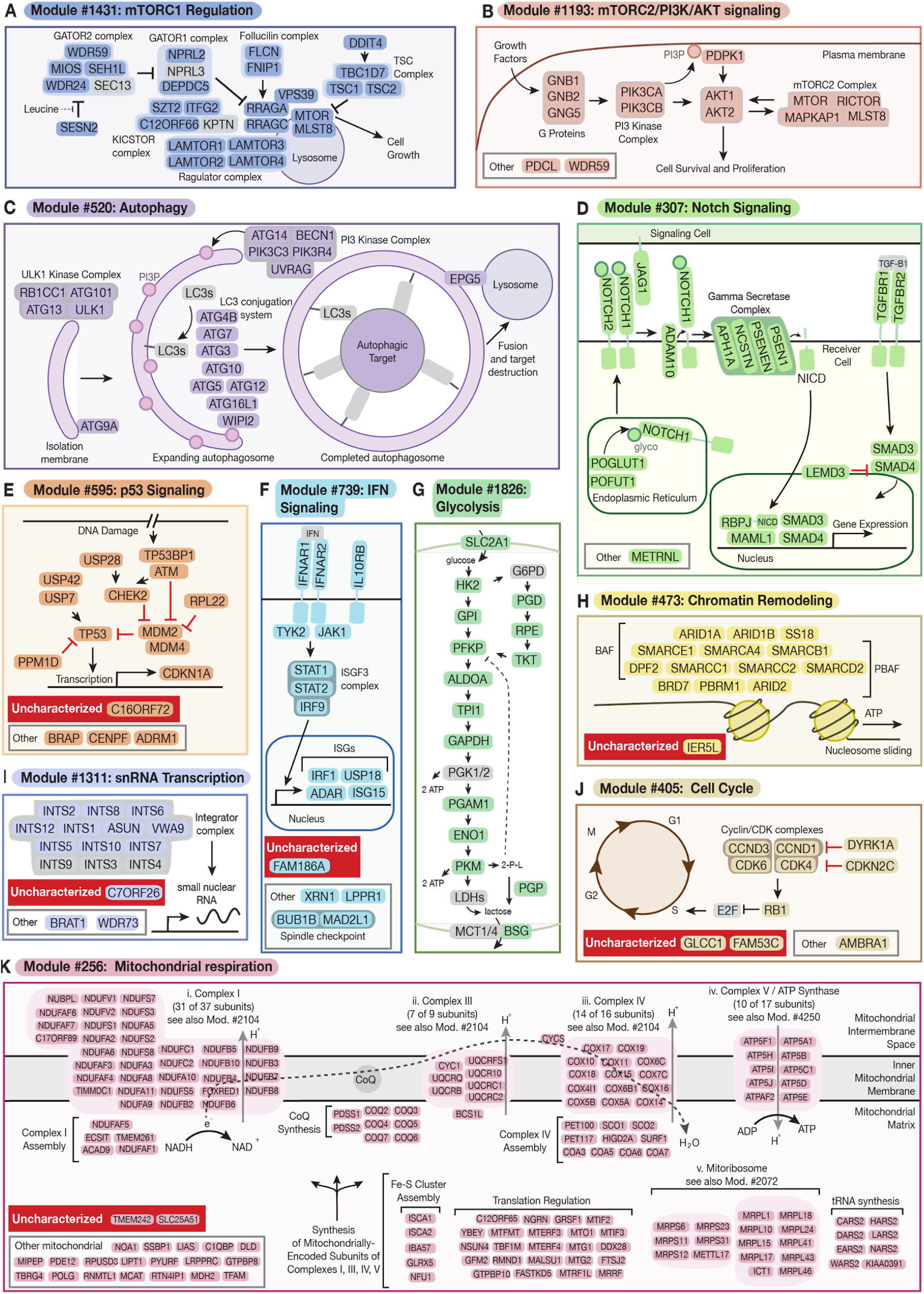
Co-essential modules recapitulate known pathways and nominate novel pathway members. (**A-K**) 10 examples of co-essential modules; all genes in each module are shown. Genes without previous evidence of pathway involvement are indicated as either “uncharacterized” (Uniprot annotation score <3) or “other”. Red inhibitory arrows between gene pairs indicate both negative regulation and negatively correlated essentiality profiles. In (A), (C), (G), (I), and (J), core pathway members not included in the module are shown in gray. In (K), subunit counts for mitochondrial respiration complexes were based on HGNC gene sets as of Oct 2019 (Povey et al., 2001). Abbreviations: (B, C) PI3P, phosphatide-inositol-3-phosphate; (C) LC3s, Microtubule-associated 1A/1B-light chain (LC3) family members; (D) NICD, Notch intracellular domain; glyco, fucose and glucose modifications transferred to NOTCH1 by POFUT1 and POGLUT1; TGF-B1, transforming growth factor beta 1; (F) IFN, interferon; ISGs, interferon-stimulated genes; (G) 2-P-L, 2-phospho-lactate (toxic byproduct of PKM) (Collard et al., 2016); (H) BAF, BRG- or HBRM-associated factors complex; PBAF, polybromo BAF complex; (K) Mod., module; CoQ, coenzyme Q.

Several important features of the co-essential modules are highlighted by the examples shown in Figure 3. First, the ability of ClusterONE to include genes in multiple modules enabled identification of pleiotropic gene functions, as illustrated by the identification of two modules containing *MTOR* that closely correspond to the two mTOR-containing complexes, mTORC1 (Figure 3A) and mTORC2 (Figure 3B) (Saxton and Sabatini, 2017). Second, co-essential modules are not limited to physical complexes, as illustrated by the near-complete identification of the glycolysis pathway (Figure 3G), or even to cell-autonomous pathways, as illustrated by the identification of the Jagged-Notch intercellular signaling pathway (Figure 3D). Third, by examining modules identified at different values of *d*, we were able to detect multiple scales of biological organization, as illustrated by the set of modules we identified that correspond to mitochondrial respiration (Figure 3Ki-v). Module #256, a 163-member module identified at *d* = 0.2, includes most nuclear-encoded subunits of the four respiratory chain complexes required for mitochondrial ATP synthesis, as well as numerous mitochondrial tRNA synthetases, elongation factors, and components of the mitoribosome required for synthesis of the mitochondrial subunits of the mitochondrial respiratory complexes (Figure 3K). Several modules identified with *d* set to 0.9, by contrast, correspond to smaller units of functional organization, such as module #4250, a 13-member module that contains 12 subunits of the ATP synthase complex (Figure 3Kiv, **Table S3**), and module #2072, a 99-member module comprising 61 subunits of the mitochondrial ribosome and many of its associated factors (Figure 3Kv, **Table S3**). Fourth, we noted that whereas several modules are nearly “complete” representations of a biological pathway, such as module #520, which comprises most of the genes identified in recent targeted screens for autophagy regulators (Figure 3C, *cf.* Shoemaker et al., 2019), and no genes not previously implicated in autophagy, many modules highly enriched for a particular pathway also contain one or more uncharacterized genes (red boxes, Figure 3E, F, I, H, J, K).

### Using co-essential modules to systematically predict the functions of uncharacterized genes

Hundreds of human genes have not been assigned any molecular function. Co-essentiality profiling has recently been used to assign uncharacterized genes to pathways, with predictions based on the functions of the genes that have the largest Pearson correlations with the uncharacterized gene (Pan et al., 2018; Wang et al., 2017). However, it has remained unclear how broadly useful co-essentiality information is in predicting the functions of the hundreds of genes that remain uncharacterized, which likely span diverse biological processes.

Co-essential modules are in many cases highly enriched for functionally related genes, and thus enable unbiased, genome-wide prediction of uncharacterized gene function. To generate a list of functional predictions for uncharacterized genes, we first mined the UniProt database to assemble a list of uncharacterized genes, which we defined as those genes with UniProt annotation score (a heuristic measure of protein annotation content) of 2 or lower. We then enumerated all the uncharacterized genes present in modules at least 100-fold enriched for one or more GO terms, excluding GO terms with < 5 genes.

The 102 uncharacterized genes assigned putative functions by this method are included, on average, in ∼2 co-essential modules, yielding a list of 220 functional predictions (**Table S4**). We excluded uncharacterized genes in syntenic modules (i.e. modules comprising genes all located on the same chromosome) from this count, since while many syntenic modules likely represent *bona fide* co-functional gene sets, others may be confounded by residual copy number artifacts or other factors (**Methods**). Notably, several of these predictions are consistent with recent experimental information that has not yet been incorporated into the Uniprot database, including *C19orf52* in mitochondrial import (Kang et al., 2016), *C16orf59* in centriole function (Breslow et al., 2018), *TMEM261* in mitochondrial respiratory complex I (Stroud et al., 2016), and *PTAR1* in Golgi function (Blomen et al., 2015), showcasing the power of this method in assigning gene functions. To prioritize a list of functional predictions for experimental validation, we ranked modules by their maximal enrichment for a given GO term, because these predictions yield the most readily testable predictions. The top uncharacterized gene predictions (ranked by GO term enrichment) span a wide range of biological processes, including mitochondrial respiration, transcription, DNA repair, Golgi function, lipid synthesis, and endocytosis (**Table S4**).

### *TMEM189* encodes the orphan enzyme plasmanylethanolamine desaturase required for plasmalogen synthesis

We selected two genes, *TMEM189* (ranked #1) and *C15orf57* (ranked #18), for experimental validation. *TMEM189*, also known as *KUA*, encodes a 270 amino acid transmembrane protein whose function was largely unexplored prior to our work, with previous studies focused on the observation that it can be transcribed as both an independent ORF and as a fusion with the neighboring *UBE2V1* gene. Both *TMEM189* and *TMEM189-UBE2V1* have been observed to localize to the endoplasmic reticulum (Thomson et al., 2000).

The top-ranked co-essential module containing *TMEM189*, module 2213, is highly enriched for genes required for synthesis of ether lipids (Figure 4A), which comprise a broad class of structural and signaling lipids involved in regulation of membrane fluidity and sensitivity to oxidative stress, and which account for approximately 20% of the phospholipids in human cells (Nagan and Zoeller, 2001). We noted that genes in this module appeared to be particularly essential in cell lines derived from haematological cancers (Figure 4B). Whereas several genes in this module, including *AGPS*, *FAR1*, and *GNPAT*, are specifically involved in ether lipid synthesis, some genes contained in this module, including *PCYT2* and *EPT1*, are required for both ether lipid synthesis and the synthesis of other ethanolamine-containing phospholipids. Based on this prediction, we hypothesized that *TMEM189* could play a role in lipid biosynthesis and have a specialized role in the synthesis of ether lipids.

**Figure 4:**
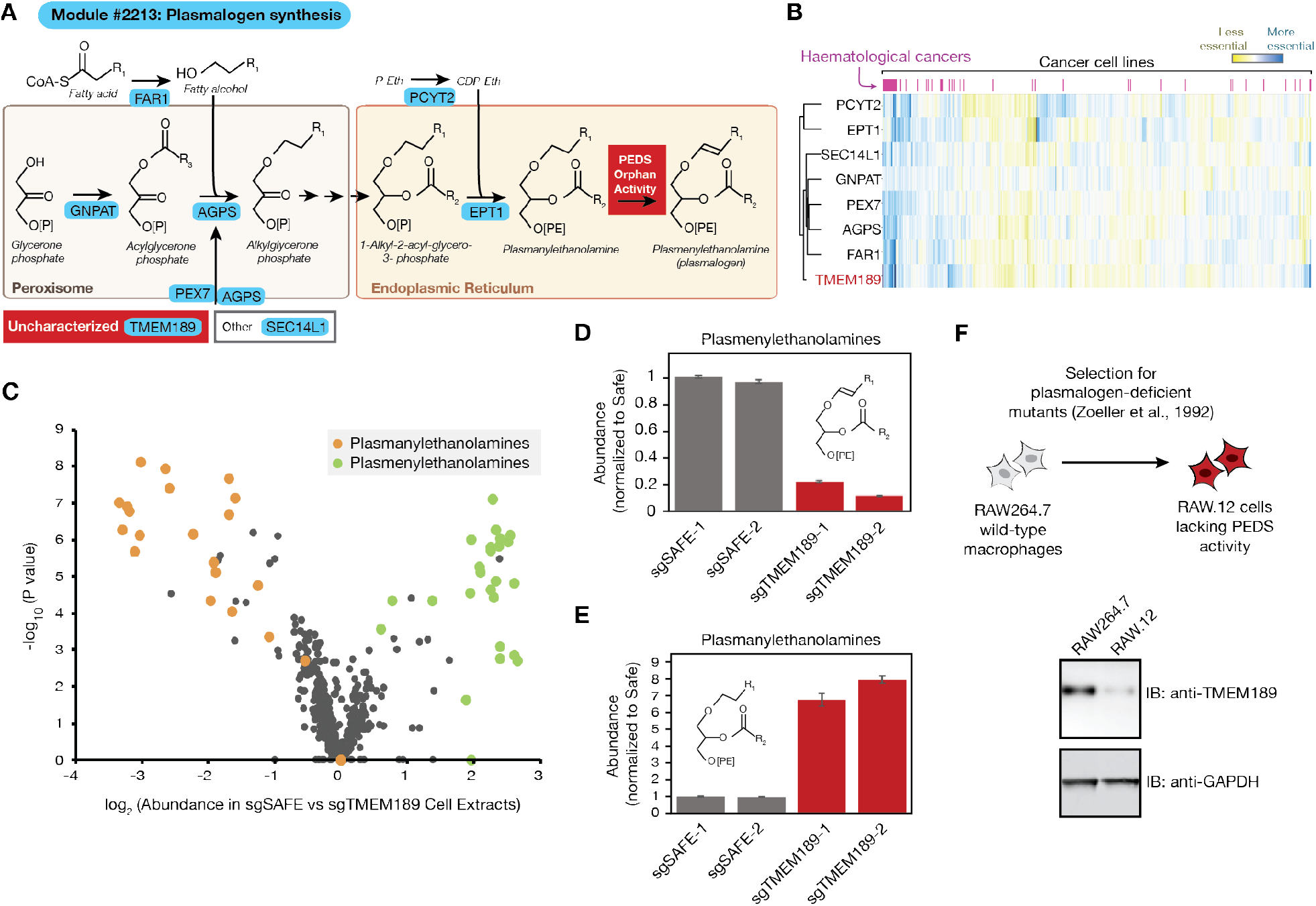
*TMEM189* encodes the orphan enzyme plasmanylethanolamine desaturase required for synthesis of plasmalogen lipids. **(A)** Schematic of module #2213 with manual annotations of gene function. Uncharacterized gene selected for validation shown in red box. PEX7 is shown importing cytosolic AGPS across the peroxisomal membrane into the peroxisome matrix (Braverman et al., 1997). Plasmanylethanolamine desaturase (PEDS) orphan enzyme activity indicated in orange. P-eth, phosphoethanolamine; CDP-Eth, cytidine diphosphate ethanolamine. **(B)** Heatmap of bias-corrected essentiality scores of genes in module 2213 in 485 cancer cell lines. **(C)** Volcano plot of all lipid species detected in lipidomic experiment, with ratio of lipid abundance in extracts derived from sgSAFE-1-expressing cells relative to sgTMEM189-1-expressing cells plotted on x-axis. **(D)** Total abundance (relative to Safe-targeting sgRNA control #1) of 37 unambiguously identified plasmenylethanolamine species in cell extracts prepared from HeLa cells transduced with indicated sgRNAs. Error bars represent standard deviation (n = 4 technical replicates). **(E)** Total abundance (relative to Safe-targeting sgRNA control #1) of 30 unambiguously identified plasmanylethanolamine species in cell extracts prepared from HeLa cells transduced with indicated sgRNAs. Error bars represent standard deviation (n = 4 technical replicates). **(F)** Top, schematic of generation of RAW.12 derivative of RAW264.7 macrophage-like line with confirmed deficiency in plasmanylethanolamine desaturase (PEDS) activity, as reported in Zoeller et al., 1992. Bottom, immunoblotting (IB) with anti-TMEM189 antibodies in RAW264.7 parental line and RAW.12 PEDS-deficient line. See also Figure S4.

To interrogate the functional role of *TMEM189* in lipid biosynthesis in an unbiased manner, we extended a targeted lipidomics method (Contrepois et al., 2018; Schüssler-Fiorenza Rose et al., 2019) to measure the absolute concentrations of several hundred lipid species in cell extracts. We compared concentrations of these lipids in cell extracts derived from HeLa-Cas9 cells that stably expressed single guide RNAs (sgRNAs) targeting either *TMEM189* or a control genomic locus. Strikingly, though the vast majority of quantified lipid species were present in similar concentrations in both types of cell extracts, cell extracts expressing *TMEM189*-targeting sgRNAs contained dramatically lower concentrations of 37 lipid species belonging to the ether lipid subclass plasmenylethanolamines (Figure 4C, 4D**, Table S5**), also known as ethanolamine plasmalogens, and higher concentrations of 30 lipid species belonging to ether lipid subclass plasmanylethanolamines (Figure 4C, 4E). Plasmanylethanolamines differ from plasmenylethanolamines in the presence of a single double bond in the sn-1 acyl chain, which forms part of the plasmalogen-defining vinyl ether bond. Plasmanylethanolamines and plasmenylethanolamines form a known precursor-product relationship, with plasmanylethanolamines rapidly converted into plasmenylethanolamines in the endoplasmic reticulum by the orphan enzyme plasmanylethanolamine desaturase (PEDS), which was first reported in mammalian cell extracts over forty years ago (reviewed in (Snyder et al., 1985)).

The accumulation of the precursors, and loss of the product, of the reaction catalyzed by PEDS in cells expressing *TMEM189*-targeting sgRNAs strongly implicates *TMEM189* as the gene responsible for orphan PEDS activity. Two orthogonal lines of evidence strongly support this conclusion. First, we examined a cell line, RAW.12, that was evolved to lack plasmalogens and shown to exhibit a specific defect in PEDS activity (Zoeller et al., 1992), and determined whether this cell line exhibits deficient expression of *TMEM189*. By immunoblotting for TMEM189 in cell extracts prepared from RAW.12 cells or its parent, unmutated cell line, RAW264.7, we confirmed that TMEM189 levels were decreased in PEDS-deficient RAW.12 cell extracts (Figure 4F). Second, *TMEM189* bears sequence features consistent with a function in lipid desaturation. *TMEM189* contains a histidine-rich domain conserved in most lipid desaturase enzymes, and is distantly related to the fatty acid desaturase *FAD4* in Arabidopsis (Gao et al., 2009), which introduces an unusual double bond in the sn-2 fatty acid (Gao et al., 2009).

We noted that *TMEM189* is also present in a co-essential module, module #808, which is highly enriched for genes involved in the biosynthesis of sphingolipids, a distinct class of lipids that predominantly localizes to the plasma membrane and, similarly to ether lipids, contributes to both signaling and membrane structure and fluidity. In lipidomic analyses, cell extracts derived from cells expressing sgRNAs targeting *SPTLC2*, a subunit of serine palmitoyltransferase, the rate limiting enzyme in sphingolipid biosynthesis, were highly depleted of several sphingolipid species, whereas abundances of most sphingolipid species were largely unaltered in cell extracts from HeLa cells expressing *TMEM189*-targeting sgRNAs, ruling out a central role for *TMEM189* in sphingolipid biosynthesis (**Table S5**). However, we observed that the relative abundances of several very long chain sphingomyelin species were altered in cells lacking *TMEM189*, with sphingomyelins with C26 fatty-acids decreased in abundance and C22 and C24 sphingomyelins increased in abundance (Figure S4A, **Table S5**). We additionally found that affinity-purified TMEM189-GFP complexes, isolated from HeLa cells, were highly enriched for SPTLC2 (Figure S4B, **Table S6**). Further work is required to determine whether this pair of observations – that TMEM189 and SPTLC2 appear to physically interact, and that the abundances of very long chain sphingomyelin species are subtly altered in *TMEM189*-knockout cells – reflects a direct role for *TMEM189* in the regulation of fatty acid incorporation into ceramides. Alternatively, because sphingolipid composition is tightly regulated to maintain membrane fluidity (Breslow and Weissman, 2010), the altered sphingolipid profile observed in *TMEM189-*knockout cells may reflect a compensatory response to loss of plasmalogens and resulting disrupted membrane composition in *TMEM189-*knockout cells. Regardless of these possibilities, our results provide conclusive evidence for a primary role for *TMEM189* as the orphan desaturase required for the final step of plasmalogen biosynthesis and provide a striking example of the power of co-essential modules to predict gene function.

### *C15orf57* is a novel regulator of clathrin-mediated endocytosis

*C15orf57* (also known as coiled-coil domain containing 32 (*CCDC32*) encodes a 185-residue protein with no annotated function. Recent reports described the existence of a chimeric transcript of unknown significance that contains *C15orf57* and the gene *CBX3* in certain tumor samples (Xu et al., 2014). *C15orf57* is present in several overlapping co-essential modules (**Table S3**), including a module (#2067) that is highly enriched for genes required for clathrin-mediated endocytosis, in particular subunits of the adaptor protein 2 (AP2) complex (Figure 5A, B). One of the best-described functions of the AP2 complex is to mediate endocytosis of transferrin bound to the transferrin receptor (Motley et al., 2003), so we hypothesized that *C15orf57* might be required for cellular uptake of transferrin. To test this, we monitored uptake of transferrin, labeled with a pH-sensitive fluorescent dye, pHrodo, by HeLa-Cas9 cells expressing sgRNAs targeting either *C15orf57*, the transferrin receptor (*TFRC*), or a control locus. Cells expressing sgRNAs targeting either *C15orf57* or *TRFC* exhibited reduced uptake of transferrin compared to cells expressing control sgRNAs, consistent with a role for *C15orf57* in transferrin uptake (Figure 5C).

**Figure 5:**
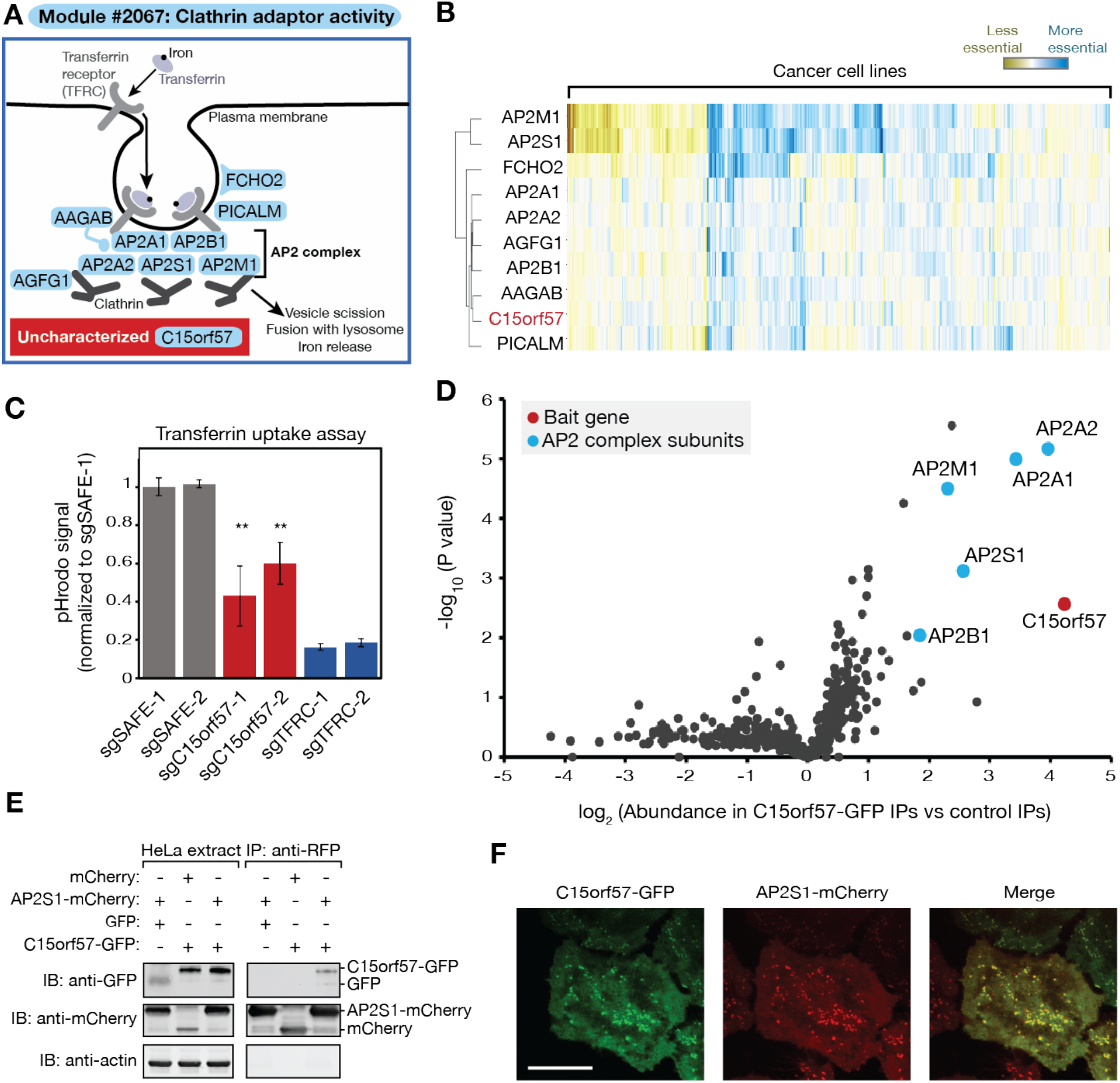
*C15orf57* is required for efficient clathrin-mediated endocytosis of transferrin. **(A)** Schematic of module #2067. Uncharacterized gene selected for validation shown in red. AP2, adaptor protein 2. **(B)** Heatmap of bias-corrected essentiality scores of genes in module #2067 in 485 cancer cell lines. **(C)** Transferrin-pHrodo uptake assay for clathrin-mediated endocytosis (24h timepoint). Error bars represent standard deviation (n = 3 technical replicates, two-tailed Student’s t-test, **p<.01). Data shown are representative of three independent experiments. **(D)** Volcano plot of mass spectrometric (TMT) analysis of C15orf57-GFP immunoprecipitates (IPs). **(E)** Extracts prepared from indicated HeLa cell extracts were subjected to immunoprecipitation with anti-RFP magnetic resin. Extracts and IP samples were resolved by SDS-PAGE and followed by immunoblotting with indicated antibodies. **(F)** Microscopy of HeLa cells transduced with C15orf57-GFP and AP2S1-mCherry constructs. Scale bar, 20µm.

To gain further insight into the mechanism by which *C15orf57* functions in clathrin-mediated endocytosis, we immunoprecipitated C15orf57-GFP complexes and analyzed them by mass spectrometry. C15orf57-GFP immunoprecipitates were strongly enriched for all five members of the AP2 clathrin adaptor complex: AP2S1, AP2A1, AP2A2, AP2M1, and AP2B1 (Figure 5D**, Table S6**). In reciprocal co-immunoprecipitation experiments, we confirmed that C15orf57-GFP physically interacts with AP2S1-mCherry (Figure 5E). We additionally confirmed through confocal microscopy that C15orf57-GFP colocalizes with AP2S1-mCherry in small puncta at the cell surface that likely correspond to clathrin-coated pits, the sites of clathrin-mediated endocytosis (Figure 5F). The identification of the members of the AP2 complex as physical interactors of C15orf57, and their colocalization in cells, suggests that C15orf57 may regulate clathrin-mediated endocytosis of transferrin (and possibly other cargoes) by directly modulating AP2 function.

### Identification of cancer type-specific pathway dependencies

A major motivation for high-throughput cancer cell line screening efforts, such as the Achilles project underlying this work, is the possibility of identifying cancer type-specific vulnerabilities that could be exploited as therapeutic targets (Tsherniak et al., 2017; Wang et al., 2017). These efforts have shown promise in identifying individual genes that are selectively essential in specific cancer types (Chan et al., 2019; Wang et al., 2017). Some cancers have also been observed to harbor selective dependencies on entire gene pathways (Hart et al., 2015, Campbell et al., 2016). We asked whether our list of co-essential modules, many of which are highly enriched for genes that function in the same pathway, could be used to identify cancer-type specific pathway dependencies.

To systematically identify differentially-essential modules across tissue types, we obtained cancer type-specific pathway dependency *p*-values for each module-cancer type pair by first calculating *p*-values for each gene and then aggregating *p*-values across genes in each module. To obtain uninflated *p*-values, we again applied GLS (see Methods). Using this conservative approach, we identified 444 modules that are differentially essential in cancers derived from 16 distinct tissue types (Figure 6A**, Table S7**).

**Figure 6:**
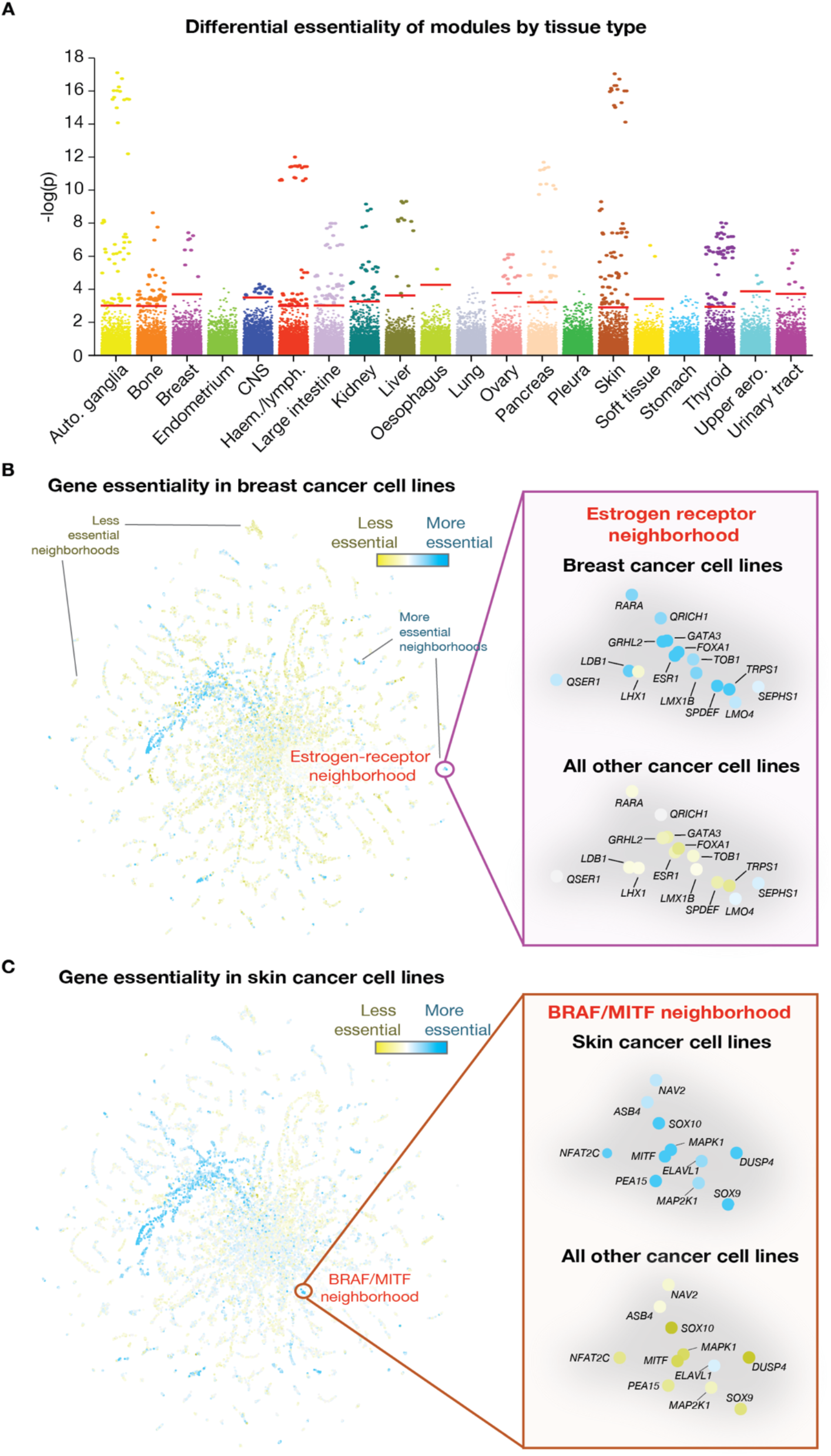
Identification of cancer type-specific module dependencies. **(A)** Differential essentiality of co-essential modules in cell lines derived from 20 tissue types. - log_10_(*p*-values) for each module are plotted for each tissue (see Methods). Red bars indicate FDR thresholds for each tissue type. Auto., autonomic; CNS, central nervous system; Haem., haematological; lymph., lymphoma; aero., aerodigestive. **(B)** Average bias-corrected gene essentiality in breast cancer cell lines plotted on two-dimensional co-essentiality network, with gene neighborhood containing *ESR1* highlighted on the right. **(C)** Average bias-corrected gene essentiality in skin cancer cell lines plotted on two-dimensional co-essentiality network, with gene neighborhood containing *BRAF* and *MITF* highlighted on the right. See also Figure S5, Video S1.

Several of the modules that are most differentially essential in specific tissue types correspond to canonical tissue-specific cancer drivers, demonstrating the power of this approach to uncover *bona fide* selective pathway dependencies. As one example, the most significantly breast cancer-specific module dependency contains *ESR1*, the estrogen receptor (ER), which is overexpressed in over 70% of breast cancers and enables hormone-dependent growth (Ariazi et al., 2006). This module (as well as the neighborhood that corresponds most closely to this module in the two-dimensional representation of the network, Figure 6B) also contains several genes that functionally interact with *ESR1*, including *SPDEF, FOXA1, and GATA3*, three master regulators of estrogen-receptor-dependent gene expression in breast cancer (Fletcher et al., 2013); retinoic acid receptor alpha (*RARA)*, a target of *ESR1-*dependent transcriptional activity (Roman et al., 1993); and *TOB1*, a gene required for estrogen-independent growth of ER-positive breast cancers (Zhang et al., 2016).

As a second example, the most significantly differentially-essential module in skin cancer (and its corresponding neighborhood) (Figure 6C) includes several components of the *BRAF*-*MAPK* pathway, which is consistent with the fact that *BRAF* is mutated in ∼50% of melanomas (Ascierto et al., 2012), as well as *MITF*, a melanoma-specific oncogene (Garraway et al., 2005) activated downstream of *BRAF*. Additional module members, including *NFATC2*, *SOX9*, and *SOX10*, have well-established roles in melanoma (Harris et al., 2010; Perotti et al., 2016). In both of these examples, the co-essential modules we identified as selectively required in certain cancer types contain sets of lineage-specific cancer drivers that are known to functionally interact, illustrating the power of our approach in identifying cancer pathway dependencies. The additional 442 modules that we identify as selectively essential in 16 cancer types (**Table S7**) represent a resource for identifying novel pathway targets in specific cancer types.

### An interactive resource for biological discovery

We created a web tool (coessentiality.net) (Figure S5; **Video S1**) to enable dynamic visualization and exploration of the genome-wide co-essentiality map shown in Figure 1C, for which we anticipate several distinct uses. First, this tool can be used as a starting point for gaining insight into the function of any gene: users can search for a gene of interest, for example, KRAS (Figure S5, **Video S1**), which is then highlighted on the interactive 2D layout in the context of its neighborhood. To understand relationships between sets of genes in a given neighborhood, users can then directly select gene neighborhoods with the cursor (**Video S1**). Once a gene set is selected, two data panels are generated: a biclustered heatmap of the genes’ essentiality profiles across the 485 cell lines and a plot of the gene set’s enrichment for annotated pathways, complexes and gene ontology terms (Ashburner et al., 2000; The Gene Ontology Consortium, 2017) (Figure S5, **Video S1**). Users can compare the essentiality profile of the selected gene set with the mutational status and expression level of other selected genes. For example, with the *KRAS*-containing neighborhood selected, users can plot the lines in which *KRAS* is mutated, which reveals that *KRAS* and several genes in its neighborhood are selectively essential in *KRAS-* mutated lines (Figure S5, **Video S1)**. Users can also gain insight into pathways that are particularly required for the growth of individual cancer lines or for cancers derived from a certain tissue by selecting cell lines (for example, U937 cells, **Video S1**) or tissue types (for example, kidney cancers, Figure S5, **Video S1**) from drop-down menus, causing the two-dimensional network to be colored according to each gene’s essentiality in the selected cell line or tissue. Finally, users can upload a specified set of genes – for example, the members of the endocytosis module containing *C15orf57* (Figure 5A**, Video S1**) – to understand relationships between multiple genes of interest. We anticipate this tool will be a broadly useful starting point for the functional characterization of genes and gene sets as well as a powerful hypothesis-generating platform for users interested in identifying cancer-type specific pathway dependencies.

## DISCUSSION

Building a global map of biological pathways in human cells and assigning function to the thousands of poorly characterized genes remain key challenges in cell biology. In this work, we demonstrate that mapping co-essentiality across a diverse spectrum of cancer cell lines enables significant progress toward both objectives.

The co-essential network developed here represents, to our knowledge, the most comprehensive and statistically robust genome-wide perturbational pathway map of human cells to date. Unlike double-perturbation approaches, our approach is scalable to all pairs of genes in the genome; and unlike prior approaches to co-essentiality mapping, it is statistically well-calibrated despite the lack of independence among the screens it was derived from. A recent comparison of modules derived from different biological networks suggested that modules created based on co-expression data are better able to recall gene relationships than co-essentiality data, in a GWAS-based benchmarking approach (Choobdar et al., 2019). By contrast, we find that co-essentiality-derived networks outperform co-expression-derived networks in their ability to recall protein complexes. The gene-gene relationships evidenced by these different datasets may be complementary, with co-essentiality especially well-powered to detect protein complexes and co-expression better able to detect certain indirect pathway relationships (Figure 2A). Our global interaction map and associated web tool showcase the high resolution and versatility of co-essentiality as a method for *de novo* pathway mapping.

Our validations of the role of *TMEM189* in plasmalogen biosynthesis and *C15orf57* in clathrin-mediated endocytosis highlight the utility of biological hypothesis generation from co-essential modules. Of note, during the preparation of this manuscript, an entirely orthogonal approach based on the study of the bacterial protein CarF, a homolog of *TMEM189*, revealed that this enzyme is responsible for PEDS activity in bacterial cells, and this activity was shown to be conserved in human cells (Gallego-García et al., 2019). The complementary approaches and orthogonal validations of *TMEM189* as the key enzyme for plasmalogen synthesis will potentiate dissection of the functions of this largely understudied class of lipids. The specific function of the plasmalogen-defining vinyl ether bond, which has been proposed to be critical for antioxidant and oxygen-sensing functions of plasmalogens, has remained difficult to assess experimentally. With the identity of plasmanylethanolamine desaturase now in hand, these and other basic questions about plasmalogen function can be addressed. Plasmalogens have been noted to be highly upregulated in a variety of malignancies, and inhibitors of this pathway have recently been explored as anti-cancer agents (Piano et al., 2015). With the discovery of *TMEM189* as a novel enzyme required for plasmalogen synthesis, we uncover an additional therapeutically targetable node in this pathway.

Our identification of *C15orf57* as a regulator of clathrin-mediated endocytosis adds another key player to this pathway; further work is required to uncover its precise mechanistic function. Nonetheless, the role of *C15orf57* in binding the AP2 complex and regulating endocytosis that we describe here may advance understanding of the significance of recurrent *C15orf57-CBX3* gene fusions that have been proposed to contribute to hepatocellular carcinoma (Zhu et al., 2019). In addition to the two uncharacterized genes for which we experimentally validated their predicted functions, we note that several additional functional predictions generated by our method are supported by evidence from other unbiased, high-throughput approaches. For example, *C7orf26*, which we predict is involved in the function of the integrator complex that is required for transcription of small non-coding RNAs (Chen and Wagner, 2010) was observed to interact with several subunits of the integrator complex in high-throughput IP-MS experiments; its expression is also highly correlated with several integrator subunits (Okamura et al., 2015; Szklarczyk et al., 2017). As a second example, the functionally uncharacterized gene *TMEM242*, for which we predict a function in mitochondrial respiration, was reported to interact with the gene product of *NDUFA3*, a subunit of mitochondrial complex I, in a high-throughput study (Szklarczyk et al., 2017). Overall, our experimental validation of two uncharacterized gene predictions, paired with our list of 100 additional uncharacterized genes for which we predict a function, provides an immediately useful resource for the broader cell biology community.

Beyond nominating functions for entirely uncharacterized genes, the modules identified in this study have the potential to suggest novel roles for genes with existing functional annotations. Within the modules that were the focus of experimental validations in this study, we note that *SEC14L1*, previously characterized for its role in inhibiting the anti-viral RIG-1 pathway (Li et al., 2013), is now the only gene in 8-gene module #2213 that has not been shown to be required for ether lipid synthesis. Notably, *SEC14L1* contains a conserved lipid-binding domain, and is related to a yeast gene, *SEC14*, involved in non-vesicular lipid transport (Saito et al., 2007). Plasmalogens are known to traffic to the cell surface after being synthesized in the endoplasmic reticulum, but the factors that regulate plasmalogen trafficking, and the route plasmalogens take to the plasma membrane, remain undefined; the possibility that *SEC14L1* regulates either plasmalogen synthesis or transport is thus a prime example of an experimentally testable hypothesis motivated by our findings. Indeed, preliminary support for this hypothesis is provided by *in vitro* studies of yeast Sec14, which revealed that purified Sec14 is sufficient to catalyze ether lipid transport between lipid membranes (Szolderits et al., 1991). Further study is required to confirm that human *SEC14L1* can similarly drive ether lipid transport between membranes and to discern whether non-vesciular lipid transport by *SEC14L1* could mediate ether lipid trafficking to the cell surface in living cells.

An additional key feature of the co-essential modules we identify, by virtue of their overlapping nature, is their ability to recapitulate multiple levels of biological organization as well as relationships between distinct pathways and complexes, as exemplified by the set of modules corresponding to distinct complexes of the mitochondrial respiratory chain (Figure 3K). The set of modules enriched for the 10-subunit endoplasmic reticulum (ER) membrane complex (EMC) provides an additional striking example. The EMC complex was recently shown to function as a transmembrane insertase required for the biogenesis of a subset of transmembrane proteins, particularly tail-anchored and polytopic proteins (Guna et al., 2018, Shurtleff et al., 2018) and is additionally noted for its role in cholesterol homeostasis (Volkmar et al., 2019). We identify several modules enriched for EMC subunits, including a 7-gene module (#5037) containing 6 EMC subunits and *TMEM147*, a gene recently shown to cooperate with EMC in transmembrane protein biogenesis (Talbot et al., 2019), and a 16-gene module (#2450) containing 8 EMC subunits and 2 genes (*MBTPS1* and *SCAP*) required for cholesterol homeostasis. In addition, we identify a 23-gene module (#534) that contains 5 EMC subunits and 12 subunits of the lysosomal V-ATPase complex (8 of which are separately contained in 9-gene module 2450). Several of these V-ATPase subunits were recently identified in an unbiased proteomic study as among the proteins most dependent on EMC function for their stability (Tian et al., 2019). Thus, the set of modules we identify that are enriched for EMC subunits correspond to known inter-pathway interactions, and demonstrate the power of co-essential modules to not only identify individual pathways but to point to possible inter-pathway relationships.

Numerous additional modules (Figure 3B, D, E, F, J, K**, Table S3**) that are highly enriched for one GO term contain components of additional pathways not previously linked to the most-enriched pathway. As one example, two essential components of the mitotic spindle checkpoint, *BUB1B* and *MAD2L1*, are present not only in a module (#1360) highly enriched for genes involved in kinetochore/spindle checkpoint function but also, unexpectedly, in a module (#739) highly enriched for interferon (IFN) response genes (Figure 3F). *BUB1B* and *MAD2L1* have well-defined roles in the prevention of chromosome instability (CIN) (Ricke et al., 2008) but have not previously been linked to interferon gene function. One possible connection is suggested by the recent observation that tumor cells with high levels of CIN generate cytosolic DNA that activates an IFN response and drives cancer progression (Bakhoum et al., 2018). *BUB1B* and *MAD2L1* prevent a particular form of CIN - aneuploidy resulting from premature sister chromatid separation – but the relevance of this type of CIN in the pathogenic process described in that study was not addressed and may warrant further investigation.

A key future direction in expanding the ability of this resource to detect functional genetic relationships is to measure additional phenotypes beyond cancer cell line growth under standard conditions. The 485 cell lines screened thus far are derived from a wide range of tissue types and exhibit a highly diverse set of mutational backgrounds, and the Achilles project plans to extend this screening to several thousand cell lines. Nonetheless, our approach has the potential to benefit greatly from screens performed in primary tissues; across individuals; under non-ambient conditions, such as in the presence of a drug or cellular stress; or with readouts other than cellular fitness, such as changes to cell morphology, gene expression, or cellular activity. Such screens offer the potential to uncover an even broader spectrum of functional interactions, and could enable a dynamic map of pathway rewiring across conditions. Overall, our genome-wide mapping of the human co-essential network comprises a powerful resource for biological hypothesis generation and discovery.

## ACKNOWLEDGMENTS

We thank Raphael Zoeller for providing RAW.12 cells and the parent cell line. We gratefully acknowledge Evan Boyle, Justin Donnelly, Mackenzie Pearson, Grace Anderson, Scott Simpkins, Trey Ideker, and members of the Bassik and Kundaje labs for helpful discussions. This work was supported by an NIH Director’s New Innovator award (1DP2HD084069-01) to M.C.B., a grant from NIH/ENCODE (5UM1HG009436-02) to A.K. and M.C.B., a Stanford Bio-X Bowes Fellowship to M.W., a Stanford School of Medicine Dean’s Postdoctoral Fellowship to R.A.K., and a Jane Coffin Childs Postdoctoral Fellowship to R.A.K..

## METHODS

### Code availability

Code to generate co-essential gene pairs, co-essential modules, modules with cancer type-specific dependencies, and the two-dimensional layout will be made available at https://github.com/kundajelab/coessentiality.

### Dataset

The dataset used to determine co-essential interactions consists of the 485 genome-wide CRISPR screens from the Achilles project 18Q3 release (Tsherniak et al., 2017). Specifically, 17,634 genes were screened in 485 cell lines from 27 distinct lineages using the Avana CRISPR library (Doench et al., 2016), and gene-level effects were quantified using the CERES algorithm to account for variability in guide effectiveness and copy number across lines (Meyers et al., 2017), resulting in a 17,634 x 485 matrix of normalized gene-level effects. Intuitively, gene-level effects represent the number of times fewer cells with the knockout doubled during the screen, compared to control cells. This dataset is publicly available at https://ndownloader.figshare.com/files/12704099, or at https://depmap.org/portal/download/all/ under release “DepMap Public 18Q3” and file “gene_effect.csv”.

### Bias correction

Bias correction was applied as described in Boyle et al., 2018. Specifically, the first 4 principal components of the gene-by-cell-line essentiality matrix across all olfactory receptor genes, defined here as those with the “olfactory receptor activity” gene ontology (GO) term (Ashburner et al., 2000; The Gene Ontology Consortium, 2017), were subtracted from the original CERES score matrix, resulting in a new bias-corrected matrix. To avoid multicollinearity and allow inversion of the covariance matrix for generalized least squares (see below), subtraction of the first 4 principal components was followed by removal of 4 cell lines (arbitrarily chosen to be the last 4), resulting in a 17,634-by-481 matrix of bias-corrected CERES scores.

### Quantifying co-essential gene pairs

The co-essentiality between each pair of genes was quantified using generalized least squares (Aitkin, 1935). In a departure from previous approaches to co-essentiality profiling, GLS automatically and flexibly accounts for the non-independence of cell lines by incorporating information about the covariation between every pair of screens. When all screens are independent and have the same variance in effect sizes across genes, the GLS effect size becomes exactly equivalent to the Pearson correlation coefficient. GLS is closely related to the linear mixed models (LMMs) used for population structure correction in genome-wide association studies (Yu et al., 2006), an analogous problem to ours.

Specifically, GLS estimates the vector of parameters **β** of the linear regression model **Y** = **Xβ** + **ε**, where **Y** is a vector of observations, **X** is a matrix of features corresponding to those observations, and **ε** are the errors or residuals, under the assumption that the mean of the errors is 0 and their variance is **Σ**, where **Σ** is a covariance matrix specified by the practitioner. The only difference from ordinary least squares (OLS) is the value of **Σ**; OLS assumes that it is the identity matrix, while GLS allows it to be any user-specified value. Here, we set **Σ** to be the covariance matrix of the data itself, i.e. V_i,j_ is the covariance of cell lines i and j across all genes in the CRISPR screen.

In practice, GLS is solved by a) inverting **Σ**, in our implementation (statsmodels.regression.linear_model.GLS from the *statsmodels* Python package) by using the Moore-Penrose pseudoinverse instead of the true inverse as a computational optimization, b) taking the Cholesky decomposition of this inverse covariance matrix **chol(Σ^-1^)**, c) transforming both **Y** and **X** by **chol(Σ^-1^)** to obtain the transformed observations **Y’ = chol(Σ^-1^) Y** and transformed features **X’** = **chol(Σ^-1^) X**, and d) running OLS on **Y’** and **X’**. (When **Σ** is the identity matrix, **chol(Σ^-1^)** is as well, so **Y’** = **Y** and **X’** = **X** and GLS reduces to OLS.)

GLS was run separately on each gene pair, resulting in a 17,634-by-17,634 matrix of GLS *p*-values. Specifically, the *endog* argument of statsmodels.regression.linear_model.GLS (the output) was set to the length-481 vector of bias-corrected CERES scores for one of the two genes, the *exog* argument (input) set to a 481-by-2 matrix where the first column is the other gene’s bias-corrected CERES scores and the second column is a constant vector of all ones (i.e. the intercept), and the *sigma* argument set to the 481-by-481 covariance matrix of the bias-corrected CERES scores. Given these three pieces of data, the GLS outputs a *p*-value indicating the statistical significance of the degree of co-essentiality between the pair of genes. Note that while the GLS *p*-value is consistent regardless of which of the two genes is chosen as *endog* and which as *exog*, the GLS effect size is not consistent with respect to this choice, and as a result is not reported. For benchmarking, GLS was also run on the non-bias-corrected data using the exact same procedure, but using the full 485 cell lines.

As a computational optimization, the rate-limiting step of the GLS calculation (inverting the covariance matrix and then taking the Cholesky decomposition) was cached and reused for each pair of genes, since all gene pairs use the same covariance matrix. With this optimization, the amortized time complexity of GLS is equivalent to Pearson correlation. The same GLS implementation was used to calculate the Pearson correlation (with and without bias correction) between each pair of genes, by setting the covariance matrix to the identity matrix.

### Pearson correlation simulations

For the simulations in Figures S1A and B, the x coordinates of the 8 red data points were sampled to be uniformly distributed between −4 and 0. The y coordinates were then sampled from 0.9 x + Normal(0, (1 - 0.9^2^)^1/2^) to have a Pearson correlation of approximately 0.9. To be visually pleasing, points were repeatedly re-simulated until two constraints were satisfied: the most extreme x and y coordinates had to be between 0.15 and 0.4 from the edge of the interval [-4, 0], and the minimum x and y differences between each pair of points had to be at least 0.2.

A second set of blue points were added alongside the red points. The blue points and red points share the same x coordinates, but the blue points’ y coordinates were sampled to be uniformly distributed between −1 and 0 to avoid having any significant correlation with the x coordinates. To enforce this lack of correlation, the y coordinates were repeatedly sampled until both the Pearson and Spearman correlation *p*-values were greater than 0.5.

In the right half of Figure S1B, the same red and blue points were plotted, in addition to 20 duplicates of each of these points, shifted by a small amount of noise sampled from Normal(0, 0.1).

### Benchmarking on CORUM, hu.MAP and STRING

For the benchmarking in Figure 3, we compared five methods: co-essentiality with GLS or Pearson and with or without bias correction, and co-expression with COXPRESdb. We used the same versions of COXPRESdb benchmarked in Pan et al., downloaded from the supplemental data to that paper at https://ndownloader.figshare.com/files/10975364 and remapped from Entrez IDs to gene names using the mapping at https://ndownloader.figshare.com/files/9120082. When benchmarking, we considered only the N = 15,552 genes present in both the Avana library and COXPRESdb.

For STRING, we used all the gene pairs in version 10.5 restricted to Homo sapiens (https://stringdb-static.org/download/protein.links.detailed.v10.5/9606.protein.links.detailed.v10.5.txt.gz). To avoid circularity, we removed gene pairs supported only by co-expression, i.e. for which the only non-zero score was for co-expression.

Following the strategy of Pan et al., we compared methods by considering their rankings on a per-gene basis. Specifically, we considered only the top N partners for each gene for N from 1 to 10, and looked at how enrichment varied as a function of N. We used the same versions of CORUM and hu.MAP benchmarked in Pan et al.

Enrichments were calculated as the percent of the top N gene pairs in the pathway or complex database, divided by the percent of gene pairs found in the database. For instance, to calculate the enrichment of COXPRESdb in CORUM for N = 2, we found the top 2 co-expressed partners per gene according to COXPRESdb (N = 2 * 15,552 gene pairs), computed the percent of these pairs that were part of the same CORUM complex, and divided by the percent of the 15,552 * 15,552 gene pairs that were part of the same CORUM complex.

Note that Boyle et al. perform an additional transformation of *p*-values after PCA correction based on the empirical null distribution of *p*-values for olfactory genes, but since this transformation is monotonic it does not affect the rankings of partner genes used in our benchmarking.

### Co-essential modules

Co-essential modules were ascertained with the ClusterONE algorithm (Nepusz et al., 2012). Briefly, ClusterONE generates modules by greedily adding nodes (genes) starting from a randomly selected seed node, so long as the sum of the edge weights within the module is sufficiently high relative to the sum of the boundary edge weights between genes in the module and their neighbors. It then merges sufficiently overlapping modules as a post-processing step, while allowing genes to be members of multiple modules (protein complexes or pathways).

ClusterONE was run on the 17,634-by-17,634 matrix of GLS *p*-values after row-wise false discovery rate correction, with edge weights set to one minus the false discovery rate q-value (Storey and Tibshirani, 2003). Default settings were used for ClusterONE, except for changing the module density parameter -d (also known as --min-density) from its default of 0.3, as discussed in the main text. For the list of modules in Table S3, all modules generated with values of d set to 0.2, 0.5, and 0.9 were merged into a single list. 11 modules that were identical at different values of d were retained in this list but were excluded from the reported count of total modules.

We noted that the resulting list of co-essential modules contained many modules that are highly enriched for genes that localize close to one another in the genome. In several cases, these modules correspond to clusters of functionally related genes that are known to colocalize in the genome, such as histone- and protocadherin-encoding genes, though in the majority of cases it remains unclear whether the presence of colocalized genes in a module reflects their shared function in a biological pathway or if it relates to vulnerabilities of CRISPR screening to copy-number artifacts that are difficult to account for perfectly (Meyers et al., 2017). Supporting the idea that co-essentiality for colocalized genes may represent a mix of true- and false-positive signals, we find substantial enrichment of syntenic gene pairs (both genes on the same chromosome) in CORUM, hu.MAP and STRING, but less enrichment than for non-syntenic gene pairs (Figure S6). We note that even after excluding syntenic modules (i.e. those that contain genes which are all located on the same chromosome), our set of co-essential modules still assigns putative functions to approximately 10,000 genes using the metric described earlier in relation to Figure 2B (i.e.the number of genes included in a module that is at least 100-fold enriched for some GO term), approximately twice as many as the next-best method (Figure S7). To enable full utilization of the dataset as well as easy discernment of syntenic and non-syntenic gene pairs and modules, we report all co-essential gene pairs and modules in Tables S1 (co-essential pairs), S3 (co-essential modules) and S4 (uncharacterized gene predictions) and annotate each as syntenic or non-syntenic.

### Identification of cancer type-specific pathway dependencies

Cancer type-specific pathway dependency *p*-values for each module and cancer type (Table S7) were obtained by 1) computing *p*-values for each gene and cancer type, then 2) aggregating *p*-values across genes in each module. In step 1), GLS was run separately for each gene with the same covariance matrix and output/*endog* argument (bias-corrected essentiality for a particular gene) as before (see “Quantifying co-essential gene pairs”). However, unlike before, the *exog* argument (input) was set to a 481-by-21 matrix of binary indicator variables for the 20 cancer types listed in Figure 6A (1 if a cell line is from that cancer type, 0 otherwise) plus an all-ones intercept column. The two other cancer types with CRISPR screen data from DepMap, cervical and biliary, were excluded due to only having a single cell line each. This multiple regression yielded 20 *p*-values for the gene, one per cancer type. We note that this approach is equivalent to an ANOVA, except using GLS instead of OLS.

In Step 2), *p*-value aggregation was performed separately for each module and cancer type using the Cauchy Combination Test/Aggregated Cauchy Association Test (Liu and Xie, 2019; Liu et al., 2019) with equal weights on all genes. In Python, this step can be expressed straight-forwardly as “module_p = cauchy.sf(np.tan((0.5 - gene_ps) * np.pi).mean())”, where gene_ps is a (number of module genes)-length vector of gene *p*-values for a particular cancer type, and module_*p* is the combined *p*-value for the module. Crucially, given that our gene-level *p*-values are highly correlated among genes in a module, the test is able to accommodate *p*-values from correlated tests (unlike the more commonly used Fisher’s combined *p* test, which uses a chi-squared instead of a Cauchy distribution to perform *p*-value aggregation), and we verified that the combined *p*-values were not inflated (median *p*-value = 0.56).

### Global structure of the co-essential network

The two-dimensional interaction map visualization was constructed to have two properties: (a) genes in many of the same ClusterONE modules are close together; (b) gene pairs with high GLS co-essentiality are close together. This was done by forming a graph G_CO_ from the ClusterONE modules (as above) and another G_GLS_ from the co-essentiality data, mixing the two with proportion α to form the mixed graph:

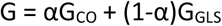

(We set α=0.99 to rely on the relatively specific and dense ClusterONE modules where possible, while falling back on pairwise GLS analysis to link genes not in any module to the rest of the network.)

The graph G_GLS_ was constructed by computing, for each pair of genes, −log(p) given by GLS between the two genes. This was denoised and compressed by keeping each gene’s edges to its 10 nearest neighbors and zeroing the other edges, resulting in each gene having a minimum of 10 neighbors in the graph. (We found our analyses fairly stable to varying the number of nearest neighbors between 4 and 100.) The graph G_CO_ was constructed using the same procedure, but with each pairwise similarity computed using the Jaccard similarity between the sets of ClusterONE modules the respective genes belonged to (for sets A and B, this is J(A,B) = |A∩B|/|A∪B|).

To visualize the network G efficiently on a global scale, we relied on the framework of *diffusion maps* (Coifman and Lafon, 2006), which basically decompose the variation in essentiality profiles over the network into short- and long-range pathway components, resulting in an embedded space for genes in the network. The genes’ positions here are relatively accurate for genes in well-separated pathways, and less so for finer distinctions – this embedded space (the “diffusion map”) is a smoothed version of the network, with each gene being represented in low dimension d = 40. The embedded space was constructed from G as follows.

G was first normalized to remove the disproportionate influence of high-degree “hub” genes in the layout, resulting in a matrix G_2_. With this gene-wise degree expanded as a matrix D_G_ = diag(∑_j_ G_ij_), the normalization operation is:

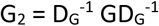

This *density normalization* further corrects for biased sampling of the network by the data (Coifman and Lafon, 2006; Haghverdi et al., 2015), as analyses on G_2_ consider the gene network corrected for the variable density of characterized genes.

The diffusion map embeds G_2_, and takes the properties of random walks on it to reveal multi-scale pathway structure. The transition probabilities of such a random walk on G_2_ are the row-sum-normalized T = D_2_^-1^ G_2_, where D_2_ = diag(∑_j_ [G_2_]_ij_).

This transition matrix T describes the evolution of any random walk, and its right eigenvectors e_1_,…, e_n_ give a diffusion map embedding when appropriately scaled. The embedding requires a parameter t, which controls the overall scale of the pathways modeled by the embedding. If the corresponding eigenvalues are λ_1_ ≥ λ_2_ ≥…, then for any t > 0, the embedded coordinates of the genes [Ф_t_]_1_, [Ф_t_]_2_, …,[Ф_t_]_40_ are:

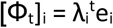

A crucial choice is that of the scale parameter t. As the current co-essentiality data are somewhat noisy for inferring fine-grained gene-gene relationships, we found it necessary to smooth them by increasing the value of t in constructing the embedding. We increased t to the minimum such that d = 40 dimensions captured 90% of the variance in the embedded space Ф_t_, and computed the resulting diffusion map Ф. This simultaneous optimization of t and Ф_t_ made the procedure adapt to and preserve large-scale global structure in a fully data-driven way, without substantive parameter tuning and using only a few matrix multiplications and one SVD.

We applied UMAP (McInnes et al., 2018) to this diffusion map embedding as in scanpy for the final global layout. Our diffusion maps implementation is in Python using the numpy and scipy packages, and includes other choices of normalization as well. The entire process ran in less than 4 minutes on the GLS- and ClusterONE-derived matrices on an Intel i7 Core CPU.

### Browser heatmap

The 481-cell-line bias-corrected CERES essentiality scores are plotted alongside the global co-essentiality network in the browser (Figure S5), and update interactively when a subset of genes is selected. The heatmap’s rows and columns are ordered by co-clustering them to find latent components, using the *sklearn.cluster.bicluster.SpectralCoclustering* implementation of the SVD-based algorithm of (Dhillon, 2001).

### GO enrichment

Below the essentiality score heatmap in the browser is the enrichment analysis panel, which displays hypergeometric *p*-values for the selected gene set against various database annotation terms, as computed by gProfiler (Raudvere et al., 2019).

### Module heatmaps

To create heatmaps for each module (Table S3), the bias-corrected CERES scores for genes in the module were hierarchically biclustered with Ward linkage using the scipy.cluster.hierarchy.linkage function from the *scipy* Python package, with the *method* argument set to ‘ward’ and the *optimal_ordering* argument set to True. This biclustering was then visualized with the seaborn.clustermap function from the *seaborn* Python package.

### Module GO term enrichments

In Table S3, GO term enrichment *p*-values were calculated via a hypergeometric test implemented using the scipy.stats.hypergeom.sf function from the *scipy* Python package. When calculating enrichments and *p*-values, genes not found in any module were excluded. GO term enrichments and *p*-values were calculated for all GO terms from the GO consortium (Ashburner et al., 2000; The Gene Ontology Consortium, 2017), except for GO terms with fewer than 20 total genes across all modules and three overly broad GO terms (biological process:biological process, cellular component:cellular component and molecular function:molecular function), which were excluded. The top 3 most-enriched GO terms for each module were listed with their hypergeometric *p*-values, provided the *p*-value was significant at a per-module Bonferroni threshold of 0.05, corrected across all GO terms.

### Generation of HeLa cell lines expressing individual sgRNAs and tagged genes

HeLa cells were maintained on tissue culture plastic and cultured in DMEM supplemented with 100 units/mL penicillin, 100 µg/mL streptomycin, and 10% fetal calf serum. Cells were passaged by incubation with Accutase, pelleting with centrifugation at 300 g for 5 min at room temperature, and replating in fresh growth medium. To transduce cells with individual constructs, HeLa cells stably expressing Cas9-BFP were lentivirally infected with constructs expressing either individual sgRNAs or GFP- or mCherry-tagged genes and a puromycin resistance (PuroR) or blasticidin resistance (BlastR) gene. At 3 d after infection, cells were selected with 2 µg/mL puromycin or 10µg/mL blasticidin for 3d, and cultured for at least 3 d without selection agent before use in experiments.

### Immunoblotting

Cleared cell extracts prepared in lysis buffer (50 mM Tris-Hcl pH 7.5, 150 mM NaCl, 1mM EDTA, 1% Triton X-100, 1 x cOmplete protease inhibitor cocktail (Roche) were heated in SDS loading buffer and subjected to SDS-PAGE, transferred to nitrocellulose, blotted and imaged using an Odyssey CLx (LI-COR Biosciences) or Supersignal West Femto Maximum Sensitivity Substrate with a Chemidoc System (Bio-Rad). The following antibodies were used: Rabbit polyclonal anti-TMEM189 (HPA059549, Sigma, 1:250), mouse monoclonal anti-GAPDH (AM4300, Fisher), rabbit polyclonal anti-mCherry (ab167453, Abcam), mouse monoclonal anti-GFP (A-11120, Thermo Fisher) and rabbit polyclonal anti-beta actin (ab8227, Abcam). In Figure 4F, the species shown is the predominant species detected using this antibody in RAW264.7 cells and HeLa cells, and corresponds to the predicted molecular weight of the *TMEM189-UBE2V1* fusion. Cell extracts from HeLa-Cas9 cells expressing sgRNAs targeting either control loci or the *TMEM189* locus (with >70% knockout efficiency verified using ICE analysis (Synthego)) were used to validate the specificity of the anti-TMEM189 antibody (data not shown).

### Time-lapse microscopy for transferrin endocytosis

HeLa cells that had been transduced with sgRNAs, co-expressed with GFP and PuroR, and selected with puromycin as described above were lifted, centrifuged, and re-plated in 24-well tissue culture plates in quadruplicate at a density of 50,000 cells per well. After 1 d, cells were washed once in dPBS, incubated in dPBS for 30 minutes, and incubated in dPBS containing 25µg/mL transferrin-pHrodo (Thermo Fisher). Plates were transferred to an incubator and imaged every 20 minutes using an Incucyte (Essen). Total red intensity for each well, averaged over 16 images per well, was calculated after applying a threshold of 1 RCU using top-hat background subtraction. Reported values represent the mean total red fluorescence intensity, normalized to the total green fluorescence signal to account for small variations in plating density, of triplicate wells. Similar results were obtained in three independent experiments using two sets of independently-generated cell lines.

### Confocal microscopy

HeLa cells were transduced with a lentiviral construct coexpressing *C15orf57*-GFP and PuroR, and then transduced with a lentiviral construct expressing A*P2S1*-mCherry and BlastR. Cells cultured in glass-bottom 24-well plates and imaged in a single plane near the glass surface using an inverted Nikon Eclipse Ti-E spinning disk confocal microscope and an Andor Ixon3 EMCCD camera using an oil-immersion 100x objective (NA=1.45). Images were assembled and adjusted for brightness and contrast in Photoshop (Adobe).

### Immunoprecipitation and mass spectrometry

For immunoprecipitations, HeLa cells that had been transduced with tagged constructs as described above were cultured in either T-150 flasks or 15 cm plates and harvested near confluency. Cell lysates for each cell line were prepared by detaching cells with trypsin, washing in PBS, resuspending in 1 mL IP buffer (50 mM HEPES, pH 6.8, 150 mM NaCl, 2mM EDTA, 1% Triton X-100, 1x cOmplete protease inhibitor cocktail (Roche)) and incubating for 30 min on ice. Cell lysates were cleared by centrifugation at 5,000 g for 5 min before incubation with 50 µl pre-washed GFP-TRAP MA beads (Chromotek) for 1 h at 4 degrees Centigrade, with end-over-end rotation. Beads were washed 4 times for 5 min with 1mL IP buffer prior to elution with 30 µl SDS sample buffer at 70 degrees Centigrade. In Figure 5E, a similar procedure was followed, except RFP-TRAP MA beads (Chromotek) were used.

For analysis by mass spectrometry, elutions were loaded on 4-12% Bis-Tris NuPage SDS-PAGE gels (Thermo Fisher) and run at 100V for 30 minutes. Gels were stained with SimplyBlue SafeStain (Thermo Fisher) and equivalent gel fragments for each lane were extracted, sliced into small fragments, and stored in 1% acetic acid. Samples were processed as described previously (Haney et al., 2018), with the following modifications. Briefly, gel slices were first resuspended in 100 μL 50 mM ammonium bicarbonate supplemented with 10 μl 50 mM dithiothreitol and incubated for 30 min at 55°C, and subsequently alkylated with 10 μl 100 mM acrylamide for 30 min at room temperature. Solution phase was discarded, and gel pieces were washed 3 times with 100 μl 50 mM ammonium bicarbonate/50% acetonitrile for 10 min at room temperature. 100 µL of 50 mM ammonium bicarbonate and 1 μg trypsin was added to digest bound proteins during an overnight incubation at 37°C. The overnight digests were spun down and the solution was collected. Peptides were extracted more two additional times with 50 µl of 70% acetonitrile/29% water/1% formic acid and incubated for 10 min at 37°C and centrifuged at 10,000 x *g* for 2 minutes, and all three extractions were combined. The combined extracts were dried using a Speedvac and reconstituted in 100 mM triethylammonium bicarbonate for TMT10plex labelling (Thermo Fisher) following the manufacturer’s instructions, and samples were mixed to generate the final peptide mixture.

Protein digests were loaded on a Waters Liquid Chromatography column coupled to an Orbitrap Fusion mass spectrometer (Thermo Fisher). Peptides were separated using a 25 cm long and 100 µm inner diameter capillary column packed with Sepax 1.8 µm C18 resin. Peptides were eluted off in a 60 min gradient at a flow rate of 600 nl/min from 5% to 35% acetonitrile in 0.1% formic acid. Mass spectrometry data was acquired by one full MS scan at 120k resolution followed with MS2 using HCD at 30k resolution. The instrument was set to run in top speed mode with 3 s cycle.

Raw data was processed using Thermo Proteome Discoverer software version 2.2. MS data were searched against a human proteome database with 1% FDR at peptide level. Protein quantification was based on the precursor ion peak intensity using the label free quantitation workflow. For Figures 5D and S4B, keratins and proteins identified with only one peptide were excluded from analysis. *P*-values were generated from Student’s t-tests between duplicate samples of indicated tagged genes and all 6 other samples analyzed in the same run (including duplicate samples derived from cells expressing GFP-tagged JTB (an unrelated gene), and from cells expressing GFP alone).

### Lipidomics

HeLa cells expressing sgRNAs targeting either safe loci or the *TMEM189* or *SPTLC2* loci were cultured in quadruplicate and harvested by centrifugation after washing with PBS. Lipids were extracted from 60 mg cell pellets using a biphasic separation with cold methyl tert-butyl ether (MTBE), methanol and water, as described previously (Schüssler-Fiorenza Rose et al., 2019). The solvent mixture contained labeled standard lipids stock (SCIEX, cat#: 5040156) to control for extraction efficiency and facilitate quantification relative to the known concentrations.

Lipid extracts were analyzed by mass spectrometry using the Lipidyzer platform (Contrepois et al., 2018), comprising a 5500 QTRAP mass spectrometer equipped with a differential mobility scan (DMS) interface (SCIEX) and high-flow LC-30AD delivery unit (Shimadzu), as described previously (Schüssler-Fiorenza Rose et al., 2019). Briefly, flow injection analysis was performed at 8 μl/min in 10mM ammonium acetate in 50:50 dichloromethane:methanol running solution, with 1-propanol included in curtain gas. DMS parameter settings were set as follows: Temperature = Low, Separation Voltage = 3.5 kV and DMS resolution = Low. PC, PE, LPC, LPE were quantified with DMS on in negative ionization mode; SM was quantified with DMS on and in positive ionization mode; FFA were quantified with DMS off and in negative ionization mode; TAG, DAG, CE, and CER were quantified with DMS off and in positive ionization mode. DMS compensation voltages were tuned using a set of lipid standards (SCIEX, cat#: 5040141), and a quick system suitability test (QSST) (SCIEX, cat#: 50407) was performed to ensure acceptable limit of detection for each lipid class. Lipid molecular species were quantified with the Lipidyzer Workflow Manager using 54 deuterated IS developed with Avanti Polar Lipids covering 10 lipid classes (SCIEX, cat#: 5040156). 17 plasmenylethanolamine species with fully saturated, 18-carbon acyl chains at the sn-1 position were excluded from analyses, as they cannot be reliably differentiated from plasmanylethanolamine species with unsaturated 18-carbon acyl chains at the sn-1 position (which are scarce in wild-type cells but are expected to accumulate in *TMEM189*-knockout cells) with the Lipidyzer platform (M. Pearson, SCIEX, personal communication).

## SUPPLEMENTAL INFORMATION

**Figure S1:**
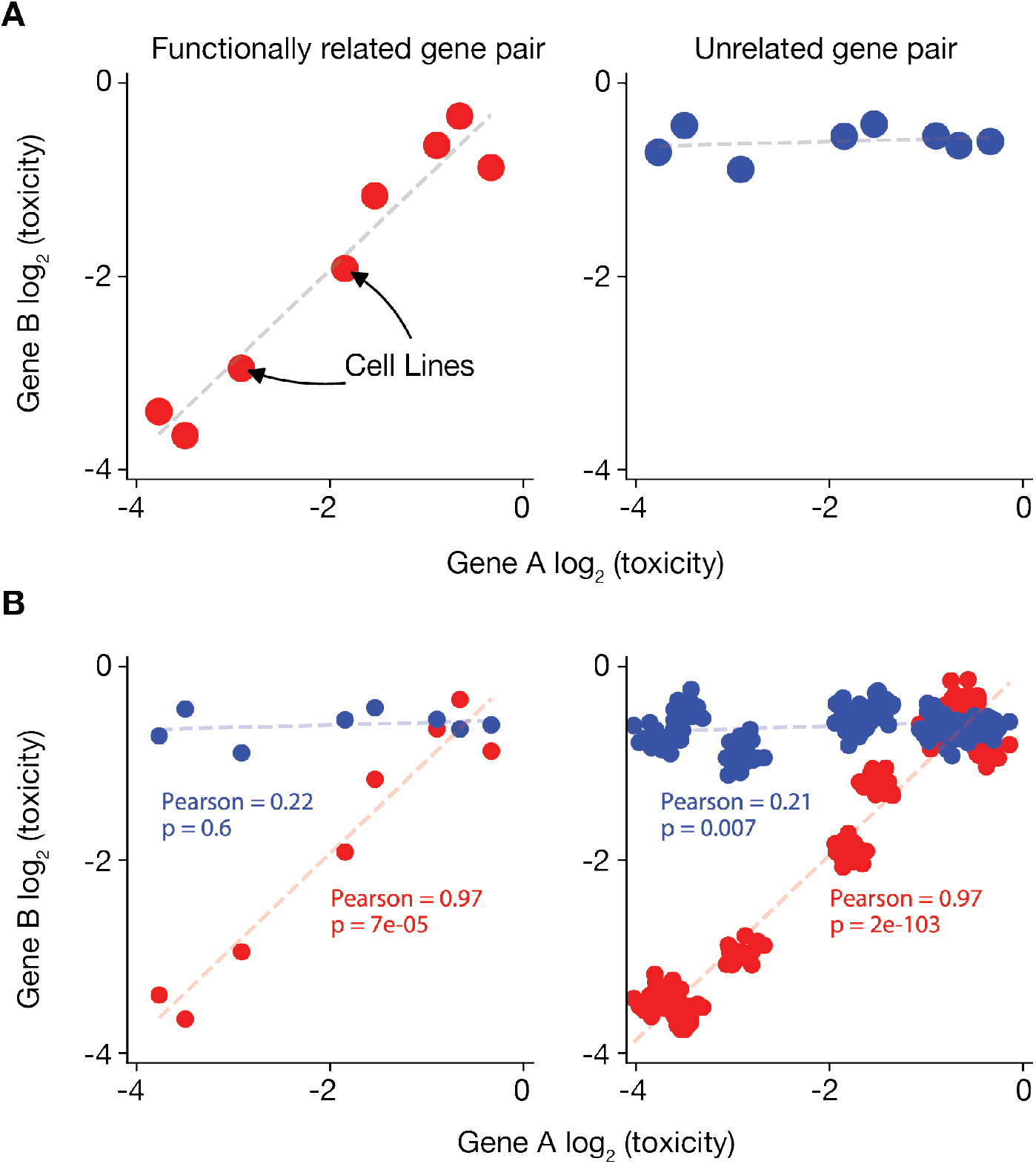
Co-essentiality profiling and the limitations of Pearson correlation. **(A)** The concept of co-essentiality: (left) a pair of functionally related genes are both essential in some cell lines and both non-essential in other lines. Essentiality can be quantified from CRISPR screens as the logarithm of the growth effect of the gene’s knockout (intuitively, the number of times fewer cells with the knockout doubled during the screen, compared to control cells). (Right) a pair of unrelated genes have uncorrelated essentiality across cell lines. **(B)** Simulation of how biological relatedness between cell lines inflates Pearson correlation *p*-values. Duplicating each point 10 times with slight noise (analogous to duplicating each screen in 10 related lines) makes the previously non-significant (p = 0.6) blue correlation highly significant (p = 0.007) and the significant red correlation (p = 7 × 10^-5^) substantially more so (p = 2 × 10^-103^), despite similar correlation magnitudes.

**Figure S2:**
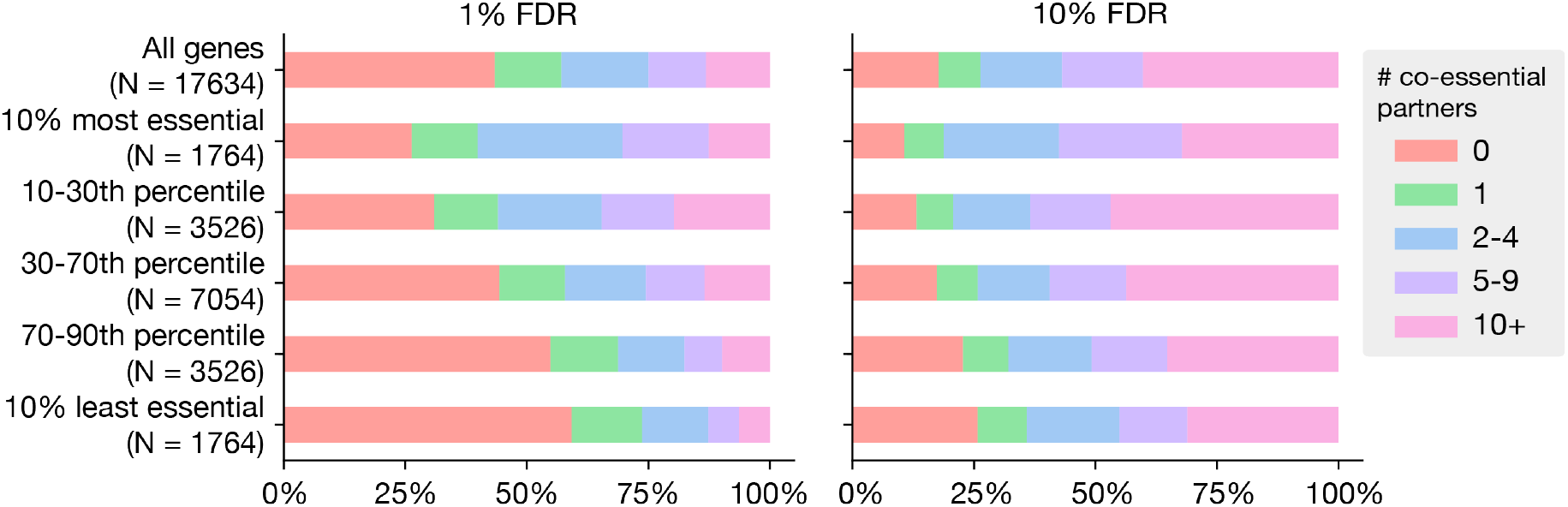
Number of co-essential partners per gene by average gene essentiality. Number of co-essential partners at 1% and 10% FDR as a function of a gene’s average essentiality (pre-bias-correction CERES score) across lines.

**Figure S3:**
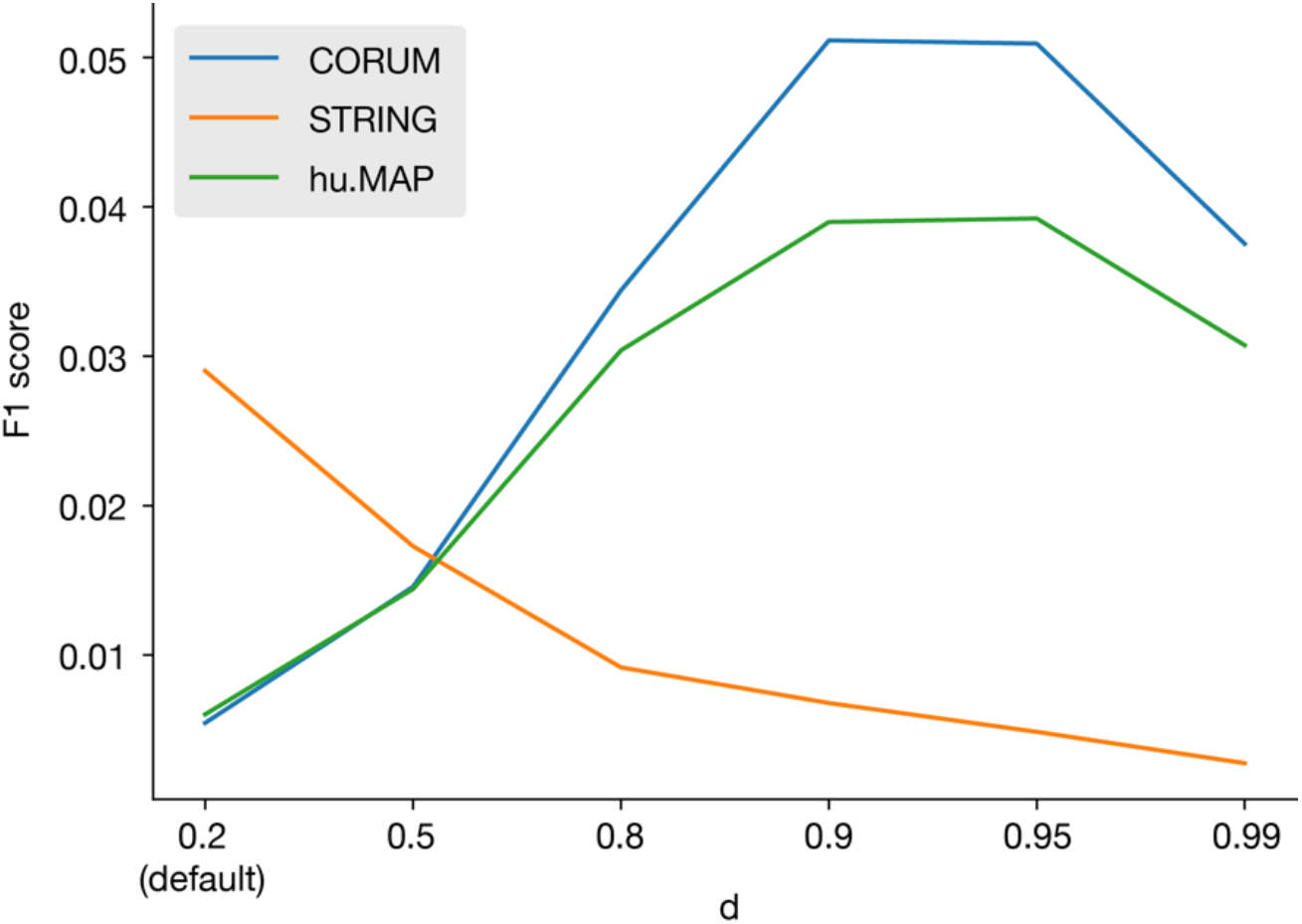
Benchmarking of cluster density *d*. F1 score (harmonic mean of precision and recall) for various values of the module density parameter *d* on CORUM, hu.MAP and STRING. F1 scores represent the performance of a binary network based on the modules (i.e. “are genes A and B in the same module?”) at predicting a binary network based on the benchmark dataset (i.e. “are genes A and B partners in the benchmark dataset?”).

**Figure S4:**
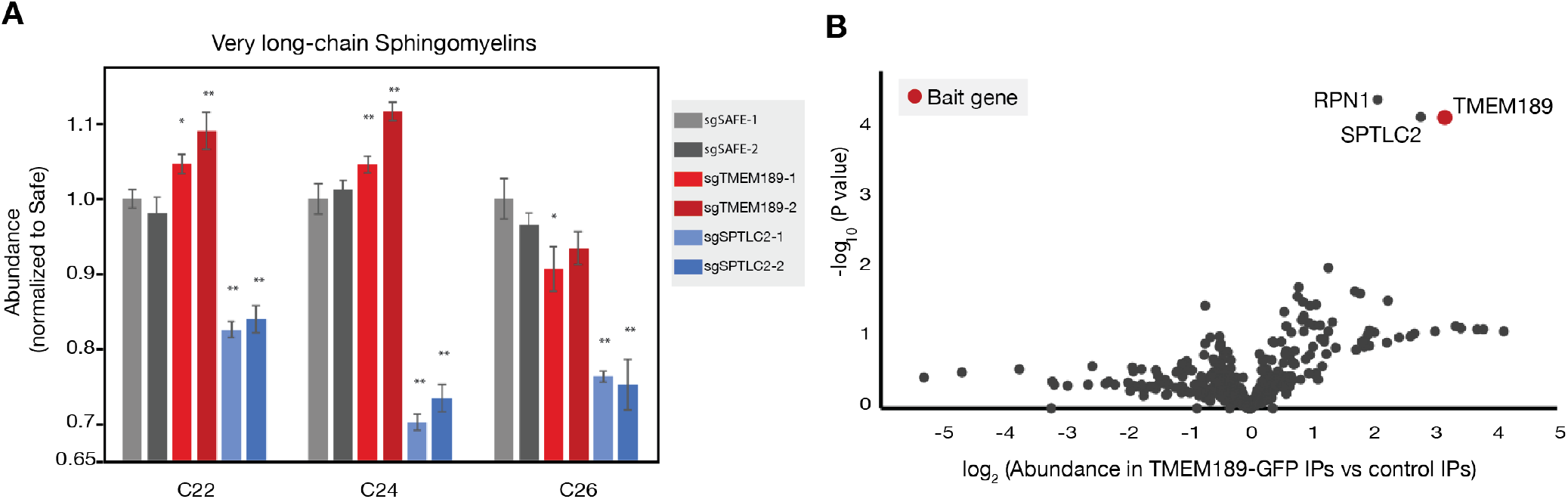
Additional functional characterization of *TMEM189* suggests a secondary role in sphingolipid biosynthesis. **(A)** Abundances (relative to Safe-targeting sgRNA control #1) of very long chain sphingomyelin species (with acyl chain length indicated on x-axis) in cell extracts prepared from HeLa cells transduced with indicated sgRNAs. sgSafe data and sgTMEM189 data are from same data set represented in Figure 4C. **(B)** Volcano plot of mass spectrometric (TMT) analysis of TMEM189-GFP immunoprecipitates. Data are from same mass spectrometry analysis as data shown in Figure 4D.

**Figure S5:**
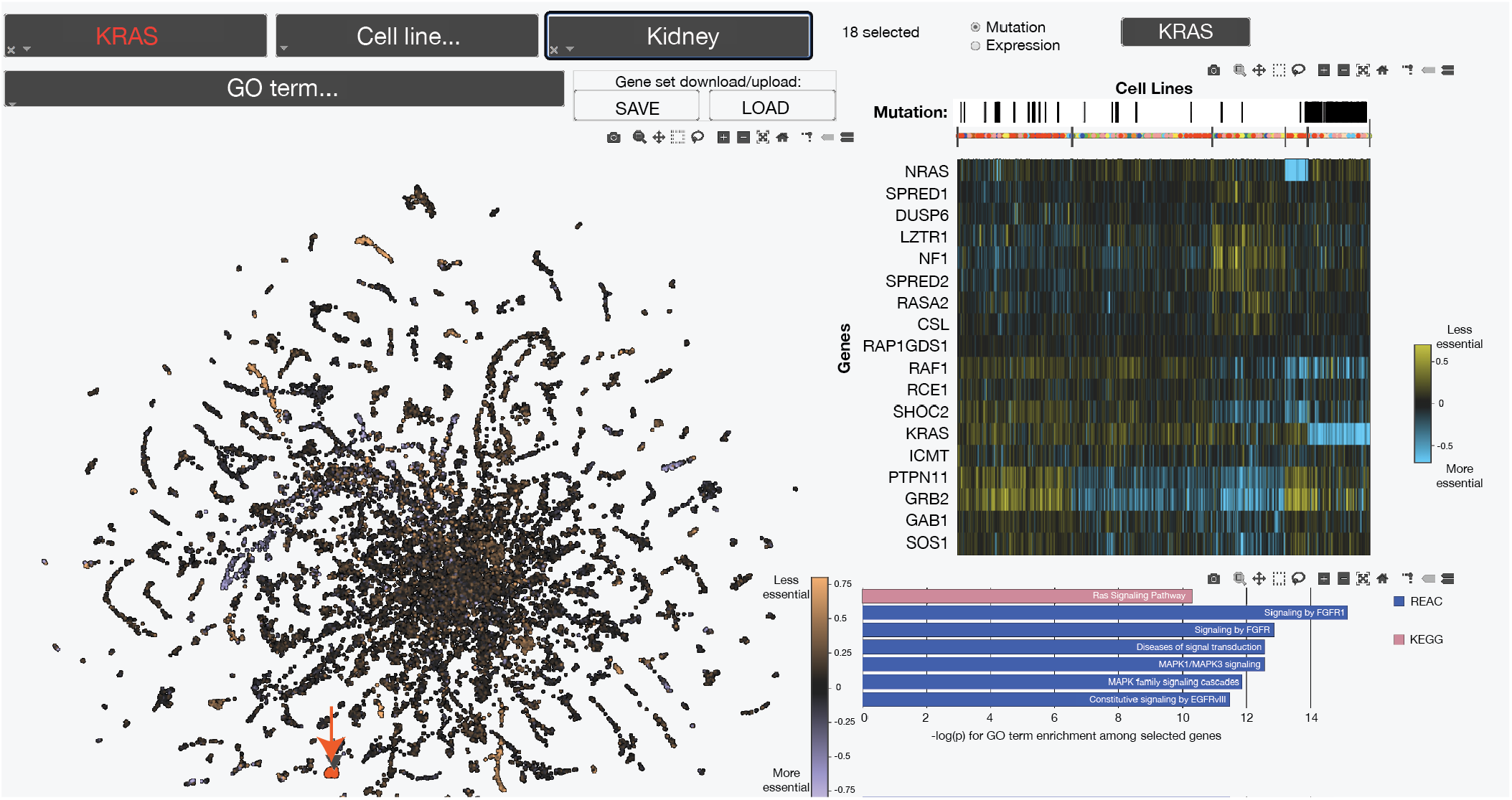
A web tool for interactive exploration of the co-essential network. Example use case for the interactive web tool (coessentiality.net). A gene, *KRAS*, was selected using the dropdown menu at top left and is marked with a red arrow in the scatterplot below. Genes selected for analysis – KRAS and its gene neighborhood – are designated with red points in the main panel (left). The heatmap panel (top right) shows that KRAS-mutant lines (selected for display using the search bar above the heat map and indicated as black marks in the “Mutation” bar above the heatmap) are enriched in a cluster (far right) that is marked by increased essentiality of KRAS. The pathway enrichment panel (bottom right) shows strong enrichments for Ras signaling and related pathways. The points in the main panel have also been selected in the tissue search bar (top middle) to be colored according to the average essentialities of each gene in kidney-derived cell lines. Gene sets can also be either saved or uploaded as csv files using the respective buttons in the top center (under “Gene set download/upload”). Some web colors and font sizes were optimized for display in this figure.

**Figure S6:**
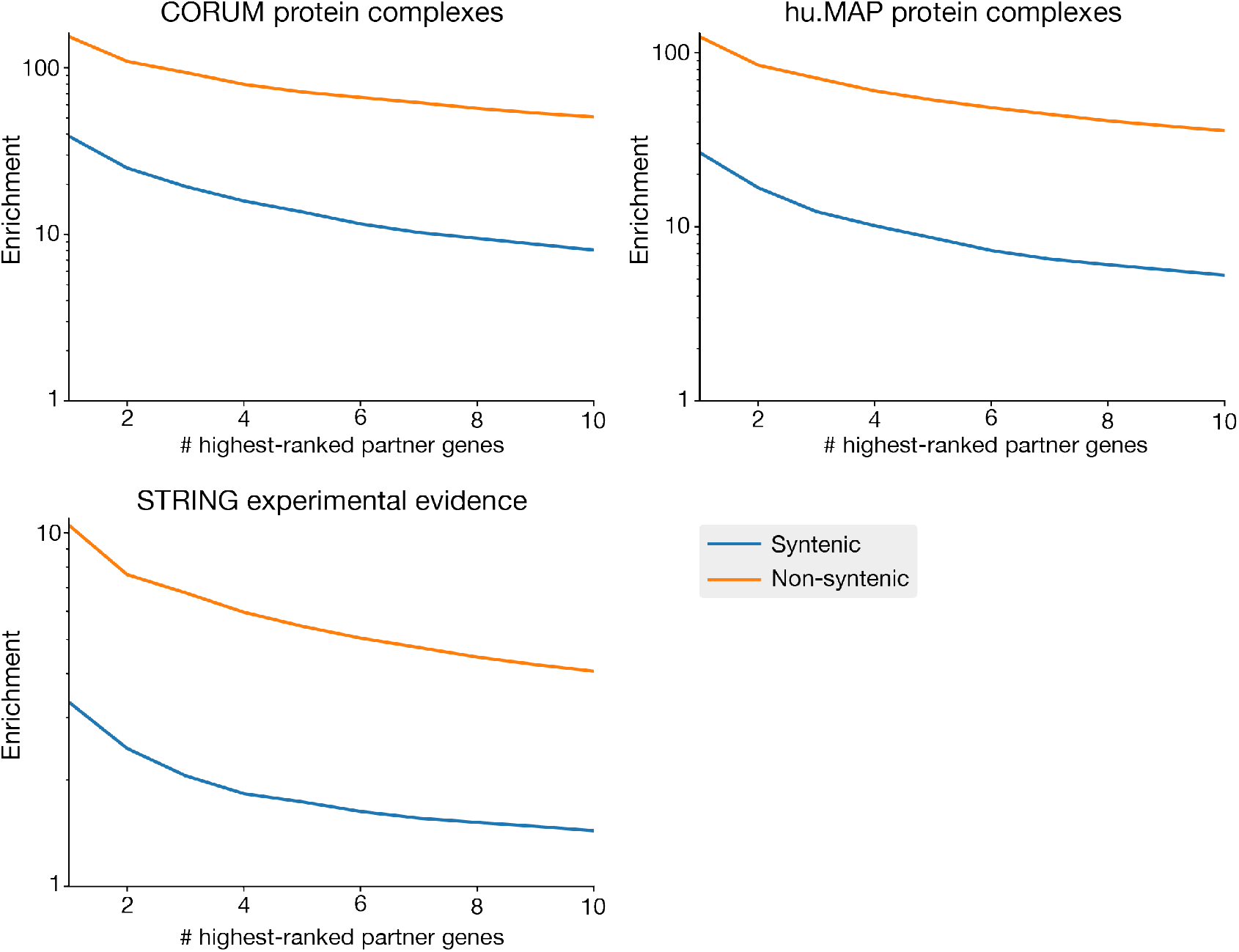
Benchmarking of syntenic versus non-syntenic modules. Enrichment of syntenic (both genes on same chromosome) and non-syntenic co-essential pairs for annotated interactions CORUM, hu.MAP and STRING databases, using the same benchmarking strategy as in Figure 2.

**Figure S7:**
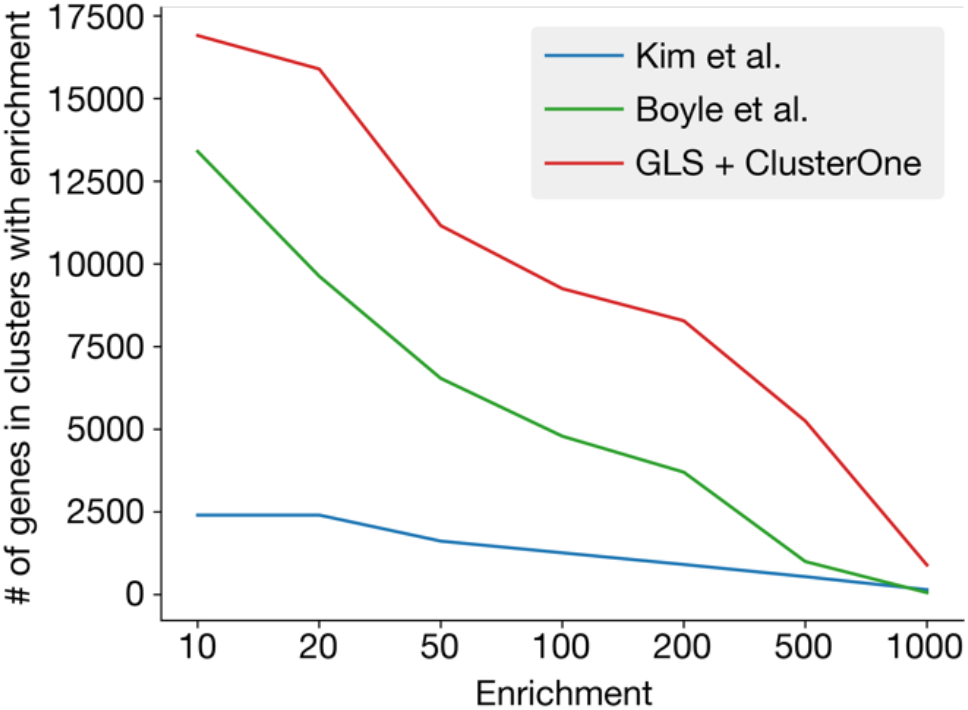
Number of genes assigned putative functions by various co-essentiality module detection methods, after excluding syntenic modules. Number of genes in non-syntenic clusters/modules at least N-fold enriched for some GO term, excluding the gene itself from the enrichment calculation, for various N from 10 to 1000.

**Table S2:**
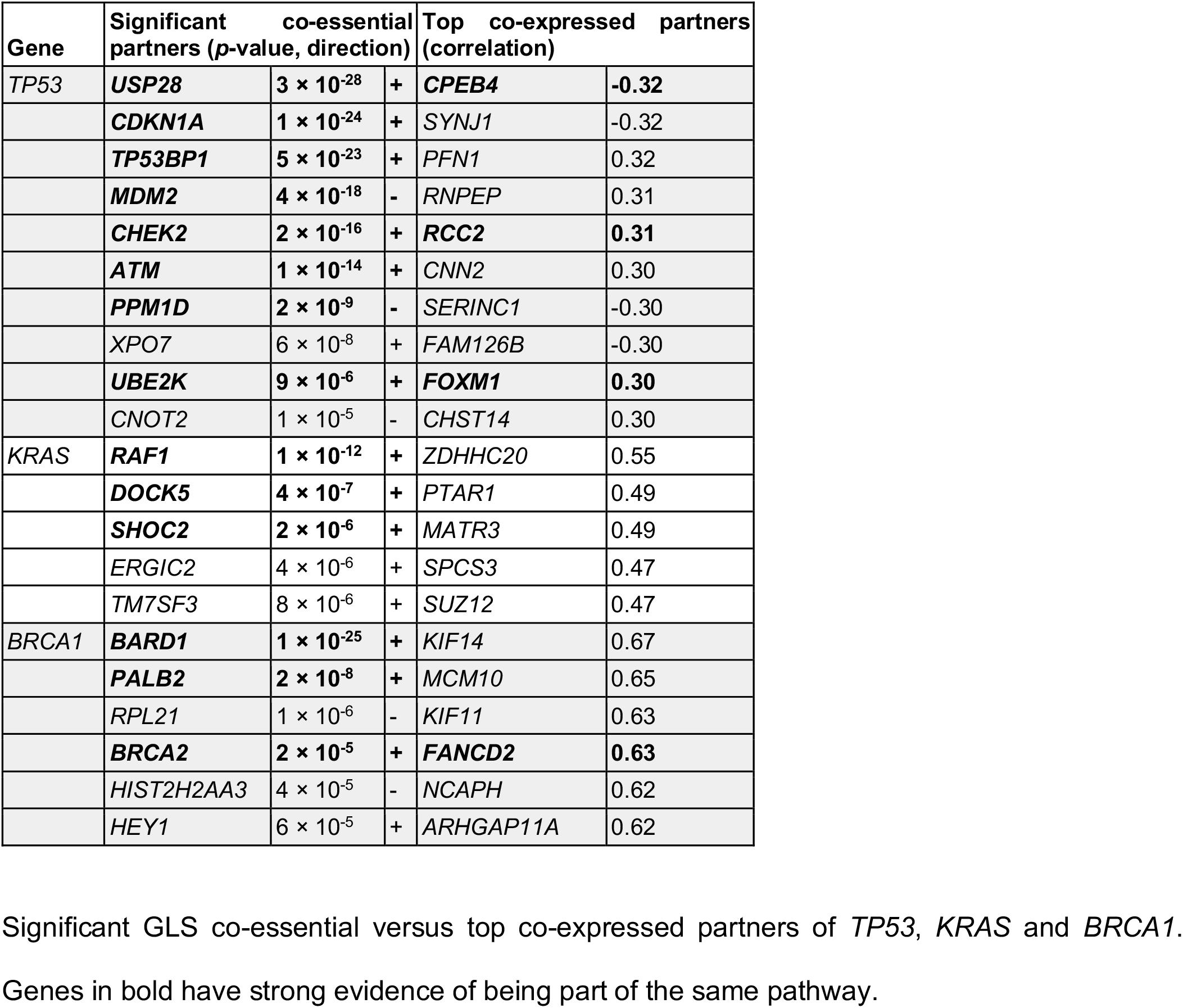
Co-essential and co-expressed partners of TP53, KRAS and BRCA1.

**Table S1: Spreadsheet of significant co-essential interactions at 10% per-gene FDR.**

List of all co-essential gene pairs identified in this study, with the number of Pubmed citations (as of Oct 2019) and chromosome location for each gene, and the direction of the gene correlation (positive (+) or negative (-)).

**Table S3: Spreadsheet of co-essential modules.**

List of all 5,228 co-essential modules and their constituent genes, with top 3 most-enriched gene ontology terms and their associated enrichments and *p-*values, the value of *d* used to define the module, and a link to the heatmap of batch-corrected essentiality data across 485 cell lines.

**Table S4: Uncharacterized gene functional predictions.**

List of uncharacterized genes that are present in co-essential modules >100-fold enriched for a gene ontology term, the Uniprot annotation score and number of Pubmed citations for each gene (as of Oct 2019), and the set of genes in each cluster that is and is not annotated with the most-enriched gene ontology term.

**Table S5: Lipidomics data.**

Lipid species concentrations for indicated lipids measured using Lipidyzer platform in indicated cell lines. QC1, QC2, and QC3 indicate quality controls (see Methods).

**Table S6: Mass spectrometry data for proteomic analysis of C15orf57 and TMEM189 interactomes.**

Proteomic data, including complete list of proteins and enrichment *p*-values, for C15orf57 and TMEM189 interactome analyses in Figures 4 and 5.

**Table S7: Cancer type-specific module dependencies.**

List of 444 differentially essential modules across 16 tissue types, ranked by *p*-value.

**Video S1: Example use cases of co-essential browser.**

Guide to use of co-essential browser showing how to navigate web tool in the context of multiple use cases, including gene lookup, gene set selection, and gene list upload.

